# Genome structural plasticity and copy number dynamics shape adaptive evolution in a barley fungal pathogen

**DOI:** 10.1101/2025.09.01.673454

**Authors:** Xiaohui Yu, Mumta Chhetri, Mafruha Hassan, Thomas Roberts, Peng Zhang, Robert F Park, Yi Ding

## Abstract

Leaf rust, caused by *Puccinia hordei*, is one of the most important diseases of barley globally. Here, we report the first haplotype-resolved chromosome-level genome assembly for *P. hordei*. To gain deeper insight and understanding of the *P. hordei* genome, we generated such assemblies for two Australian isolates with contrasting virulence, one collected in the 1970s and one in 2001. The two haplotype-phased and chromosome-level assemblies were used to investigate genome architecture in *P. hordei* and to investigate variability and evolutionary relationships of 41 virulence-divergent isolates that were collected over a 54-year period (1966–2020) in Australia. We report, for the first time in a rust fungus, evidence of a conserved chromosome, chromosome 9, which is enriched for structural variation and accessory features that imply it functions as a compartmentalized core chromosome with regions subject to rapid diversification, potentially acting as a structural hotspot in adaptive evolution in *P. hordei*. Clear genetic stratification into clonal and recombining lineages was uncovered, with evidence of both long-term asexual propagation and recombination. Analyses of haplotype-resolved mating-type gene phylogenies and chromosome 9 k-mer profiles confirm divergence at key loci underpinning reproductive modes, distinguishing clonally derived from sexually recombined groups. We also identified pervasive copy number variation across the *P. hordei* genomes, with clonal lineage-specific duplications. Consistent with this finding, the study for the first time links copy number variation of the *Cyp51* gene to fungicide insensitivity. This latter discovery is highly significant as it represents the first documented case of fungicide insensitivity in a rust pathogen in which an underlying genetic mechanism has been identified. Together, these findings illuminate how genome structure, recombination, and structural variation shape the evolution and adaptation of *P. hordei*, providing a guidance for future surveillance and management of this pathogen.

## Introduction

Barley leaf rust, caused by the basidiomycete fungus *Puccinia hordei* Otth., is one of the most important diseases threatening global barley production (Clifford, 1985; Park, 2003). Breeding barley cultivars with genetic resistance is an effective and sustainable strategy for managing this disease (Park et al., 2015). However, virulence matching deployed resistances has often emerged following the deployment of resistance (Park, 2003), with many of the designated all stage resistance (*R*) genes being rendered ineffective (Park et al., 2015). A deeper understanding of the biology and of virulence adaptation in *P. hordei* requires improved genome resources and detailed studies of its genomic content.

Like other cereal infecting rust species, urediniospores and aeciospores of *P. hordei* have two separate haploid nuclei, and it exhibits a complex macrocyclic, heteroecious life cycle that includes alternation on species of the genera *Ornithogalum*, *Leopoldia* or *Dipcadi* (Duplessis et al., 2014; Park et al., 2015). Current understanding of the biology of some cereal attacking rust species has been greatly improved by haplotype-phased genome assemblies and comparative genomics. Such studies have revealed substantial genomic plasticity in these rust pathogens, driven by transposable elements (TEs), gene duplications, and large-scale structural variations (Duplessis et al., 2014; Schwessinger et al., 2020). The ability to generate high-quality genome assemblies has further improved the resolution of complex genomic regions, enabling the characterization of mating-type loci, effector repertoires, and mechanisms underlying pathogenicity (Guan et al., 2025; Henningsen et al., 2024; Luo et al., 2024; Tam et al., 2025; Wu et al., 2021).

Many cereal rust race-specific R proteins specifically recognize pathogen-derived effector proteins, commonly referred to as avirulence (Avr) proteins. Avr recognition can trigger a localized immune response that limits pathogen growth at the infection site (Jones & Dangl, 2006). However, rust pathogens can evade host immunity by evolving loss-of-function Avrs, enabling infection of previously resistant plant varieties. Barley leaf rust *R* gene cloning has progressed significantly, with recent identification of those encoding two conventional nucleotide-binding leucine-rich repeat domain containing (NLR) type immune receptors *Rph1* and *Rph15*, the membrane executor-like *Rph3*, transcription factor *Rph7*, and the lectin-receptor kinase *Rph22* (Chen et al., 2021; Chen et al., 2023; Dinh et al., 2022; Dracatos et al., 2018; Wang et al., 2019). In contrast, no *Avr* gene has yet been isolated from *P. hordei*. Understanding *P. hordei* virulence patterns and population shift structures by building high-quality genomic resources and accurate gene annotation are key to confirming Avr candidates, which will serve as invaluable markers in future pathogen surveillance.

Population dynamics and adaptive responses to selective pressures such as fungicide application play a critical role in shaping the evolutionary trajectories of many fungal plant pathogens. In agroecosystems, extensive use of demethylation inhibitor (DMI) fungicides exerts strong selection pressure that often drives the rapid emergence of fungicide insensitive pathogen lineages (Cools & Fraaije, 2013; Lucas et al., 2015). In Australia, *P. hordei* has been documented to undergo sexual recombination on *O. umbellatum*, an introduced species that is considered an environmental weed that has become naturalized in many parts of south-eastern Australia (Wallwork et al., 1992). Long-term race analysis has documented recurrent shifts in pathotype composition and virulence acquisition, suggesting both clonal proliferation and recombination contribute to adaptation (Park, 2003). However, the genetic mechanisms underlying adaptation to fungicide pressure remain unknown. Recent work in other plant and human fungal pathogens including *Rhynchosporium commune*, *Zymoseptoria tritici*, and *Candida albicans* has shown that variations especially in genes like *Cyp51* can mediate fungicide insensitivity through gene duplications and dosage effects (Ford et al., 2015; Mair et al., 2016; Stalder et al., 2023). Pathogenicity surveys of *P. hordei* samples collected across Australia from 2021 onwards indicated that clonal lineage 5453 P+, first identified in Western Australia in 2001, exhibits insensitivity to the DMI fungicide tebuconazole. Fungicide insensitivity was further detected in other pathotypes, raising the possibility that genomic variation mechanisms may underlie azole resistance (Park et al., 2022). Elucidating the population structure and genome structural variations in *P. hordei* is thus critical for understanding its evolutionary potential and for developing sustainable disease control strategies.

Rust genomes contain extensive TE content and a higher number of predicted genes compared to other basidiomycetes, with significant intraspecific variation (Corre et al., 2025; Cuomo et al., 2017). This complexity made the development of accurate genome assemblies difficult, with early attempts generating haplotype collapsed assemblies with dramatically compromised contiguity. Chen et al. (2019) reported a preliminary genome assembly for *P. hordei* that despite being incomplete provided initial insights into gene content and especially the effector complement of this damaging pathogen. The sequencing and assembly of cereal rust genomes (Li et al., 2023; Liang et al., 2023) have been enabled by long-read technologies, particularly PacBio high fidelity (HiFi) sequencing and assemblers with sophisticated algorithms such as hifiasm and hicanu (Cheng et al., 2021; Nurk et al., 2020; Wenger et al., 2019). Haplotype phasing is crucial in analyzing heterozygous diploid rust genomes. Advances in phasing and genome-continuity improving tools such as chromosome conformation capture (Hi-C) have enhanced the assembly of haplotype-phased contigs to scaffold or chromosomal scale resolution without the need for genetic and physical maps (Kronenberg et al., 2021). To gain deeper insight and understanding of the genome of *P. hordei*, we generated the first haplotype-phased and chromosome-level genome assemblies for *P. hordei,* using two Australian isolates of contrasting virulence. The two haplotype-phased and chromosome-level assemblies were used to investigate genome architecture in *P. hordei* and to investigate variability in the pathogen from 1966–2020 in Australia. They provide a valuable genomic resource for an important plant pathogen and a crucial foundation for identifying key genes responsible for pathogen adaptation, elucidating virulence evolution, and advancing the genomic study of *P. hordei*.

## Materials and methods

### Plant and rust materials and rust inoculation

*Puccinia hordei* isolates Ph518 and Ph560 were both sourced from liquid nitrogen storage at the Plant Breeding Institute Cobbitty, The University of Sydney. Ph518 was isolated in eastern Australia in the 1970s and is avirulent for all *Rph* genes except *Rph8* and *Rph10*. Ph560 was isolated in 2001 in Western Australia, and is virulent for *Rph1*, *Rph2*, *Rph4*, *Rph6*, *Rph8*, *Rph9*, *Rph10*, and *Rph12*. It is considered to be a probable exotic incursion, and served as a founding isolate that gave rise to a series of single-step derivative mutant pathotypes (Chen et al., 2019; Park RF, unpublished). The two isolates were purified from single pustules with 2-3 rounds of increase on the susceptible barley genotype Gus. To obtain enough urediniospores for high molecular weight DNA extraction, seedlings of the barley genotype Gus treated with maleic hydrazide (0.5g/L, 50 ml per pot) at the coleoptile stage were used for the final round of increase. During inoculation, a differential set of barley genotypes that carried most characterized *Rph* leaf rust resistance genes (Park et al., 2003) was included to confirm the identity and purity of the two isolates. An additional collection of 31 *P. hordei* isolates together with eight previously reported ones (Chen et al., 2019) across a time span of over 50 years (Supplementary Table 1) were used for population genomics analyses. These isolates were purified and increased using the same method as Ph518 and Ph560. Inoculation was performed at approximately 12-14 days after sowing. Urediniospores suspended in mineral oil (Isopar L®) were sprayed onto plants using a mist atomizer. The inoculated plants were incubated in a dark humid chamber at ambient temperatures overnight and then moved to 26°C for disease development. Infection types were assessed at 8-10 days post inoculation, using a modified “0”-“4” scale as described by Park and Karakousis (2002).

### DNA extraction and sequencing

High molecular weight DNA of Ph560 and Ph518 was extracted from 1 g of urediniospores using a slightly modified CTAB extraction method and then purified as described by Yu (2024). Library preparation and HiFi long read sequencing were performed using one single-molecule real-time sequencing (SMRT) cell with the PacBio Sequel II Platform (Novogene, Singapore) for each isolate. A total of 30.9 Gb and 37.15 Gb HiFi reads with average read lengths of 17,252 bp and 18,515 bp were generated for Ph518 and Ph560, respectively. Hi-C sequencing was performed for isolate Ph560. DNA crosslinking was first carried out with an incubation of 100 mg of urediniospores in 15 ml of 1% formaldehyde solution at room temperature for 20 minutes. One gram of glycine was then added to the sample, and the mixture was incubated at room temperature for 20 minutes with occasional vortexing. The resulting sample was then centrifuged at 1,000 × g for 1 minute, followed by removal of the supernatant. The spores were then subjected to DNA extraction and Hi-C library preparation at Phase Genomics (Seattle, WA, U.S.A) using Proximo Hi-C Kit (Microbe). The libraries were sequenced at Genewiz (South Plainfield, NJ, USA) to generate 100 million 150-bp paired-end reads. For isolates used in population genomics, approximately 50mg of urediniospores were used to extract genomic DNA following Ding et al. (2021). Pair-end (150-bp) short reads library was prepared and sequenced at over 50′ coverage using the NovoSeq X platform at Novogene (Singapore). Raw reads are deposited at NCBI under BioProject: PRJNA1292008.

### Phased genome assembly and cleanup

For Ph560, the Hi-C integrated mode of hifiasm v0.19.5 was used to generate the initial haplotype-phased assembly (Cheng et al., 2021). The HiFi and Hi-C reads were fed into hifiasm for genome assembly with the parameters “-t22 -l3 -s0.55”. The resulting haplotype assemblies were blasted against a mitochondrial database retrieved from NCBI (https://ftp.ncbi.nlm.nih.gov/refseq/release/mitochondrion/) to identify and filter out mitochondrial sequences. Database creation and the subsequent sequence blasting were performed using BLAST+ as per documentation (Camacho et al., 2009), after which, the haplotype assemblies were subjected to removal of haplotig sequences using Purge Haplotigs v1.1.2 (Roach et al., 2018). First, a subset of HiFi reads was mapped to each haplotype using minimap2 v2.22 (Li, 2018) with the parameters “-ax map-hifi --secondary=no”, and the resulting alignment files were sorted and indexed using samtools (Danecek et al., 2021). Lastly, the “hist”, “cov”, and “purge” commands of Purge Haplotigs were executed sequentially to remove haplotig sequences.

### Hi-C scaffolding and telomere identification

For Ph560, Hi-C scaffolding was carried out using YaHS v1.1a (Zhou et al., 2022). Raw Hi-C reads were mapped to each haplotype assembly according to Arima Genomics mapping pipeline (https://github.com/ArimaGenomics/mapping_pipeline/blob/master/arima_mapping_pipeline.sh). The resulting alignment files and haplotype assemblies were fed into YaHS to produce initial scaffolded assemblies using command “yahs” with additional parameters “-e GATC --telo-motif TTAGGG”. The outputs were processed using “juicer pre -a” command of YaHS to generate text files of Hi-C links, which were then loaded into Juicer tools to produce Hi-C contact maps. The resulting files were visualized and curated manually using Juicebox v1.11.08 (Durand et al., 2016). The final scaffolded haplotype assemblies were generated using the “juicer post” command of YaHS. The metrics of the scaffolded haplotypes were assessed using QUAST v5.0.2 and BUSCO v5.7.1 (basidiomycota_odb10) (Gurevich et al., 2013; Simão et al., 2015).

Telomeric motifs in each scaffolded haplotype were identified using the Telomere Identification toolKit (Tidk, v0.2.64) (Brown et al., 2025). The “tidk search” command was first performed to search for telomeric repeats (TTAGGG/CCCTAA) across each genome, and the resulting occurrence counts were visualized using “tidk plot” to assess the presence of telomeres at the chromosome ends.

Hi-C data were not generated for Ph518 due to technical reasons. HiFi only mode of hifiasm with default parameters was used to generate chromosome-scale contigs, and Hi-C data from Ph560 was used to assist scaffolding. The resulting haplotype assemblies of Ph518 were subjected to homology-based misassembly correction using RagTag (Alonge et al., 2022). RagTag was executed with additional parameters “-f 5000 -d 500000 -b 10000 --inter” to only detect and break large structural misassemblies. After correction, the assemblies were then scaffolded using “scaffold” command to generate final chromosome-level haplotype assemblies. Assembly cleanup, assessment of assembly metrics and telomere identification were performed using the same methods as described earlier.

### Repeat annotation and long terminal repeat (LTR) aging

The chromosome-level haplotype assemblies of the two isolates were soft-masked before genome annotation. A *de novo* repeat library was first created for each haplotype using “LTRstruct” mode of RepeatModeler v2.0.1 (Flynn et al., 2020). Simultaneously, the haplotype assemblies were soft-masked by RepeatMasker v4.1.0 (Smit et al., 2013-2015) using the built-in repeat database of *Puccinia* with parameters “-species puccinia -xsmall”. The annotated *Puccinia*-specific repeat sequences were extracted from each haplotype using rmOut2Fasta.pl, a Perl script from RepeatMasker, and then merged with the corresponding *de novo* repeat library for the final repeat masking.

Of the outputs of repeat masking, alignment files of repeat elements in each haplotype were processed by calcDivergenceFromAlign.pl (a Perl script of RepeatMasker) to calculate divergence metrics, i.e. Kimura%. LTR retrotransposons were then extracted to enable the calculation of LTR insertion times as follows: Age (in million years) = Divergence (Kimura%) / (2 × substitution rate). A substitution rate of 2% per million years was adopted as described previously (Baranova et al., 2015).

### Gene structure annotation and effector prediction

In a previous study by Chen et al. (2019), RNAseq data (100-bp paired-end) was generated for Morex plants inoculated with the *P. hordei* isolate Ph612. The RNAseq data (BioProject: PRJNA495764) and orthologous protein sequences from other *Puccinia* species (Henningsen et al., 2022; Li et al., 2023; Li et al., 2019; Schwessinger et al., 2022) were used as transcriptome and protein evidence, respectively, for gene structure annotation. Specifically, each soft-masked haplotype was indexed using “genomeGenerate” command in STAR v2.7.10a with the parameter “--genomeSAindexNbases 12” (Dobin et al., 2013). Illumina RNAseq reads were quality checked by FastQC and trimmed by fastp (-l 50 -f 15 -t 10) (Andrews, 2010; Chen et al., 2018). Trimmed RNAseq reads were then mapped to each haplotype to generate sorted alignment files. Two independent *ab initio* gene predictions were then performed using BRAKER3, incorporating either transcriptomic or proteomic evidence separately (Gabriel et al., 2023).

Genome-guided *de novo* transcriptome assembly was conducted for each RNAseq alignment file using Trinity (Grabherr et al., 2011). The resulting transcripts were processed with the PASA pipeline, generating transcript alignment files using three different aligners: BLAT, GMAP, and minimap2 (Haas et al., 2003). Finally, the *ab initio* gene predictions and transcript alignments were integrated using EVidenceModeler to produce final gene structure annotation for each haplotype assembly (Haas et al., 2008).

Secreted proteins were predicted using either Phobius (Käll et al., 2004) or a combination of SignalP-4.1g and TMHMM-2.0c (Krogh et al., 2001; Nielsen, 2017). For the latter approach, annotated proteins from each haplotype were analysed with SignalP-4.1g for signal peptide prediction using parameters “-t euk -U 0.34 -u 0.34”. Protein sequences, excluding the predicted signal peptide regions, were then processed with TMHMM-2.0c to identify transmembrane helices. Proteins predicted to contain a signal peptide but lacking transmembrane helices were classified as secreted proteins. Among these, proteins smaller than 300 amino acids were further considered as candidate effectors.

### Haplotype genome comparisons and comparative genomics

Haplotype assemblies were aligned using minimap2 (-x asm5 -c --eqx) for pairwise comparisons or Mummer (nucmer --maxmatch -l 100 -c 1000) (Marçais et al., 2018) for haplotype comparison and structural variation detections. To visualize haplotype similarity in a pairwise manner, a heatmap was generated based on the percentage of alignments with sequence identities greater than 50% using D-GENIES (Cabanettes & Klopp, 2018). Synteny analysis and visualisation were conducted using SyRi and plotr (Goel et al., 2019). ModDotPlot v0.8 (static -id 80 -k 11 -r 3000) was used to perform sequence alignment and visualisation of chromosome 9 (Chr9) (Sweeten et al., 2024). To test statistics of Chr9 genomic content comparisons across quantile levels, the R packages GenomicRanges, rtracklayer and Biostrings were used (Lawrence et al., 2009; Lawrence et al., 2013; Pagès et al., 2025). To compare *P. hordei* with other five *Puccinia* genomes, *viz*. *P. coronata* f. sp. *avenae* (*Pca*, isolate 12SD80) (Henningsen et al., 2022), *P. graminis* f. sp. *tritici* (*Pgt*, Pgt21-0) (Li et al., 2019), *P. polysora* f. sp. *zeae* (*Ppz*, GD1913) (Liang et al., 2023), *P. striiformis* f. sp. *tritici* (*Pst*, 134 E16 A+ 17+ 33+) (Schwessinger et al., 2022) and *P. triticina* (*Pt*, Pt15) (Li et al., 2023), orthologous gene clusters were identified using the OrthoFinder algorithm within OrthVenn3 (Sun et al., 2023). Each cluster represents genes across different *Puccinia* species that are derived from a single gene in their last common ancestor. Only the first/primary haplotype assembly of each *Puccinia* species was included in the analysis, with Ph560A representing the *P. hordei* genome. Phylogenetic analysis was conducted using the Maximum likelihood method in JTT (Whelan & Goldman, 2001) + CAT (Lartillot & Philippe, 2004) model, with proteins from single-copy orthogroups serving as independent evolutionary units. Additionally, Gene Ontology (GO) enrichment of species-specific orthologous gene clusters and collinearity visualization were performed using the built-in utilities of OrthVenn3 (Sun et al., 2023).

### Hi-C and genome compartment analysis

Hi-C contact matrices were generated, normalised and visualised using runHiC v0.8.7 with default settings (Wang, 2016), and R packages HiCExperiment v0.99.9, HiCool v1.7.1, HiContacts v1.9.1 and InteractionSet v1.36.1 (Lun et al., 2016; Serizay et al., 2024). Chromosomal compartment analysis and cis-contact eigenvector calculation were conducted using the python package cooltools v0.7.1 (Open2C et al., 2024).

### Population genomics

Quality checks and trimming of Illumina whole genome sequencing reads were performed using the same methods described above. Trimmed reads were then mapped to Ph560 references with BWA (bwa mem -t8 $ref $fq1 $fq2 | samtools view -b -S -h -F 4) (Danecek et al., 2021; Li, 2013). The resulting bam files were sort-indexed then fed into FreeBayes for variant calling with default parameters (Garrison & Marth, 2012). The variant calling files generated were then filtered using vcffilter with “DP > 10 & QUAL > 10” (Danecek et al., 2011). For principal component analysis (PCA), vcf files were fed into PLINK 1.9 for linkage pruning (plink --vcf $VCF --double-id --allow-extra-chr --set-missing-var-ids @:# --indep-pairwise 50 10 0.1 --extract --make-bed -pca) (Chang et al., 2015). The PC1-PC4 variance were extracted from eigenvec and eigenval files and k-mean clustered with confidence ellipse as significance. For population structure analysis, admixture and admix were used following previously reported methods (Ding et al., 2021). We used FineSTRUCTURE to explore fine-scale population structure based on haplotype sharing (Lawson et al., 2012). Co-ancestry matrices were generated using ChromoPainter v2 with default parameters and a species-specific recombination map (Lawson et al., 2012). FineSTRUCTURE was run with 1 million burn-in and 1 million MCMC iterations, sampling every 1,000 steps. The final clustering and tree were inferred from the posterior distributions and visualized using the FineSTRUCTURE genetic clusters and co-ancestry patterns among individuals. To detect introgression between populations, we used Dsuite to compute D-statistics and the f-branch (f_b_) statistic, where the tree topology reflected prior population relationships (Malinsky et al., 2021). For phylogenomics, iqtree and ggtree were used for consensus tree building and visualisations (Minh et al., 2020; Yu et al., 2017). Decomposition tree network was constructed using the Neighbor-Net algorithm with SplitsTree v4.17.0 (Huson, 1998). Branch confidences of both phylogenomic trees and Split trees was based on 1,000 bootstrap replicates.

### Copy number variation (CNV) calling

The reference genome was divided into non-overlapping 10 kb bins. For each isolate, read depth per bin was calculated using the coverage command of BEDTools, excluding bins overlapping >50% low-complexity or repetitive regions (Quinlan & Hall, 2010). To account for differences in sequencing depth and coverage bias, raw depth values were normalized within each isolate by dividing by the genome-wide median depth. A 10 kb-window median smoothing was applied to reduce technical noise and to enhance CNV signal. Normalized and smoothed read depth data were processed using the DNAcopy algorithm implemented in the R package DNAcopy v1.70.0 (Seshan & Olshen, 2025). Circular binary segmentation was applied to each sample independently to detect statistically significant shifts in read depth, representing candidate CNV breakpoints. The segmentation algorithm used a significance threshold (alpha = 0.01) and required a minimum of three bins per segment. Segments with a log2 read depth ratio (segment mean) above +0.3 were classified as duplications, and those below –0.3 as deletions. Segments within ±0.3 were considered normal copy number. To determine gene-level copy number status, annotations were overlapped with CNV segments using BEDTools intersect. Genes were labelled as duplicated or deleted if ≥50% of their transcript length overlapped a CNV segment in the respective category. The remaining genes were considered normal copy-number. A gene was considered affected by a CNV if at least 50% of its transcript body overlapped with a CNV segment. Genes with multiple transcripts were resolved to the transcript with the most extensive overlap. To identify shared or sample-specific CNVs, gene-level CNV calls were aggregated across all samples. For each gene, the proportion of samples exhibiting deletions or duplications was computed. Genes affected by the same CNV event in over 90% of samples were considered fixed CNVs, while those present in a subset of samples were designated polymorphic CNVs. To reduce potential misclassification due to low coverage or edge effects, genes with both deletion and duplication in different samples were excluded from specific enrichment tests unless consistent evidence supported a particular CNV type. Distances between CNV-affected genes and the closest TE were calculated using the RepeatMasker-generated TE annotations. Gene coordinates were compared against TE coordinates using the GenomicRanges package in R. The shortest distance from each gene to the nearest TE was recorded, and distances were summarized per CNV category (deletion, duplication, normal). TE annotations were parsed to extract TE class and family and CNV-TE associations were visualized using ggplot2 (Wickham, 2016).

### K-mer analysis

For k-mer counting, trimmed Illumina and HiFi reads were split into k-mers using the meryl count function from the Meryl suite v1.4 (Rhie et al., 2020). K=31 was used for the purpose of extracting sequence annotation. To extract haplotype-specific signal, the Ph560 haplotype-resolved assemblies (Chr9A and Chr9B) were used as k-mer query references. Sample-by-k-mer matrices were constructed for each chromosome separately (Chr9A and Chr9B) that aggregated k-mer presence across samples. To minimize low-information content, the matrices were filtered to retain only the top 100 most variable k-mers based on variance across samples. These matrices were used to generate clustering heatmaps, revealing distinct patterns of k-mer presence in isolate groups. To reduce redundancy among highly similar k-mers and to identify sequence motifs representative of haplotype-specific sequence classes, we clustered k-mers based on sequence similarity. Pairwise Hamming distance was calculated for all k-mers, and clusters were defined by single-linkage agglomerative clustering with a maximum Hamming distance threshold of 1. Each k-mer cluster was assigned a unique ID, and for each sample, the number of k-mers per cluster was counted. This yielded a reduced matrix of sample-by-k-mer-cluster counts. These matrices were used to generate the collapsed heatmaps. To enhance interpretability, hierarchical clustering was applied to both samples and k-mer clusters. To examine whether the highly variable k-mer clusters encode sequence motifs potentially relevant to regulatory function, the top most variable clusters were used for motif enrichment using the MEME Suite v5.5.1 (Bailey et al., 2015) in DNA mode with default parameters.

### Identification of mating-type genes

To identify mating-type genes in *P. hordei*, coding sequences of orthologous mating-type genes from *Pt* and *Pst* genomes were used as queries for sequence blasting against the Ph518 and Ph560 haplotypes. In the *Pt* genome assembly Race 1 BBBD, *bW-HD1* alleles correspond to *PTTG_09683* and *PTTG_27730*, while *bE-HD2* alleles are *PTTG_10928* and *PTTG_03697*. The putative pheromone receptor (P/R) complex is represented by *PTTG_28830* (*STE3.1*), *PTTG_09751* (*STE3.2*), and *PTTG_09693* (*STE3.3*) (Cuomo et al., 2017). Similarly, in the *Pst* genome assembly (Pst-130) (Vasquez-Gross et al., 2020), genes *FUN_008986*+*FUN_008987* and *FUN_010468* were used as blasting queries for *bW-HD1* alleles, while *FUN_008988* and *FUN_010469* for *bE-HD2* alleles. The genes *FUN_000740*, *FUN_005623*, and *FUN_017677* were used to search for *STE3.1*, *STE3.2*, and *STE3.3*, respectively (Holden et al., 2023). Orthologous clustering of mating-type genes across *Puccinia* species was analyzed using OrthoFinder3 (Emms & Kelly, 2019). To validate the functional relevance of the observed k-mer divergence, we compared the k-mer-based isolate groupings with gene trees constructed from mating-type genes *STE3.2* (on Ph560 Chr9A) and *STE3.3* (on Ph560 Chr9B). Maximum likelihood trees were constructed using iqtree v2.2 with the GTR+G model and 1000 ultrafast bootstrap replicates. Trees were visualized using ggtree in R, and tips were colored by k-means cluster membership derived from the PCA analyses. Motif logos and barplots were generated using R and ggplot2. Phylogenetic trees were rendered with ggtree.

### Cloning and yeast complementation assays

To clone the *PhCyp51* gene, full length Ph560 cDNA (1633 bp) was amplified with primers listed in Supplementary Table 6 and inserted into the cloning site of the vector pYES2.1 (ThermoFisher). The generated vector was then confirmed by sequencing and introduced into the *S. cerevisiae* strains BY4743 and Δ *erg11*, respectively. For yeast growth assays, cultures of different strains were grown overnight in synthetic dropout (SD) media using 2% glucose as carbon source. Overnight cultures were thoroughly washed before resuspending with SD supplemented with 2% galactose for induction of expression. Prothioconazole at 1/10 low field rate was supplemented in the agar plates.

## Results

### Haplotype-phased genome assemblies

Two *P. hordei* isolates, Ph560 and Ph518, were selected to generate haplotype-phased, chromosome-level assemblies. For Ph560, initial haplotype assemblies were produced using hifiasm in Hi-C integrated mode, resulting in haplotype A (Ph560A) with a total length of 158.5 Mb and an N50 of 10.6 Mb and haplotype B (Ph560B) with a total length of 154.6 Mb and an N50 of 11.8 Mb. After assembly cleanup and Hi-C scaffolding, both final haplotypes contained 18 chromosome-sized scaffolds with cumulative lengths of 142.6 Mb and 144.0 Mb for Ph560A and Ph560B, respectively. The chromosomes were numbered based on synteny with other *Puccinia* genomes. To validate phasing quality, we performed Hi-C mapping on the diploid genome assembly comprising both haplotypes. The absence of excessive noise and the well-aligned chromosome boundaries indicated a high assembly contiguity and accurate Hi-C mapping (Figure 1A). The genome-wide contact matrix showed a clear diagonal of high interaction frequency, indicating strong intra-chromosomal contacts. Two major blocks along the diagonal correspond to Ph560 A and B chromosomes, respectively, with minimal signal between them (Figure 1A). A breakdown of contact types across Ph560A chromosomes (Figure 1B and Supplementary Figure 1) shows that cis interactions dominate (∼75-80%) across most chromosomes. This confirms successful haplotype phasing and spatial location of chromosomes in each nucleus.

**Figure 1:**
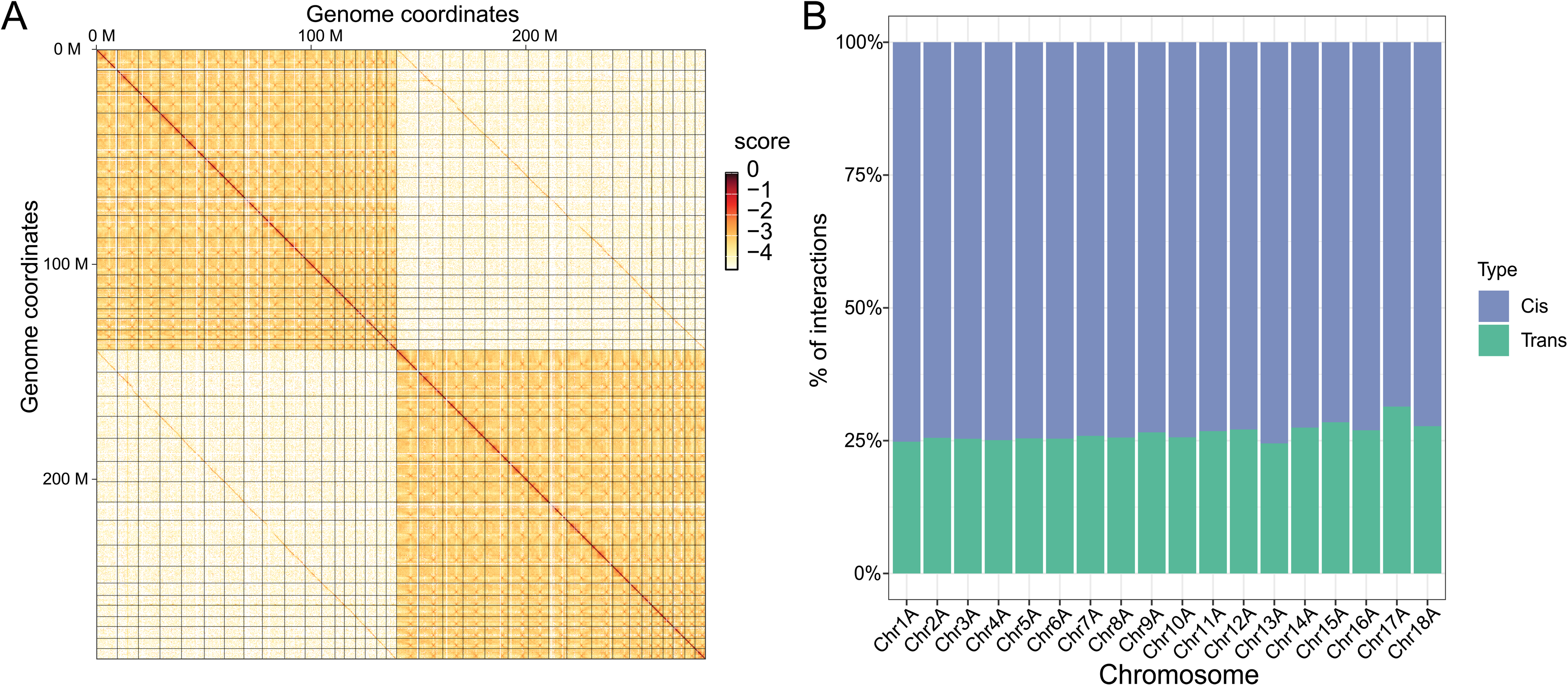
Haplotype-resolved assembly of Ph560 guided by Hi-C scaffolding. (A) Hi-C contact maps of the two Ph560 haplotypes, with haplotype A on the top left and haplotype B on the bottom right along the diagonal. Shown as balanced core with log10 scale at a resolution of 100 kb window. (B) Proportion of reads mapping to haplotype A chromosomes representing cis contacts confirms accurate haplotype phasing. Stacked barplot showing the percentage of intra-chromosomal (cis, blue) and inter-chromosomal (trans, green) Hi-C interactions for each haplotype A chromosome. The absence of excessive trans contacts further indicates minimal cross-haplotype mapping errors and high assembly contiguity.

Searches for telomeric motifs identified 35 telomeres in each haplotype (Figure 2A), with all chromosomes except chromosomes 3A and 13B assembled to telomere-to-telomere resolution. BUSCO analysis revealed 95.0% and 94.3% gene completeness for Ph560A and Ph560B, respectively (Table 1). Over 90% of assessed BUSCOs (Ph560A: 91.5%, Ph560B: 90.4%) were both complete and single-copy. BUSCO completeness for the combined Ph560 haplotypes reached 95.5%. Collectively, these new assemblies exhibited greatly improved continuity and completeness compared to the previously published *P. hordei* genome (Chen et al., 2019).

**Figure 2:**
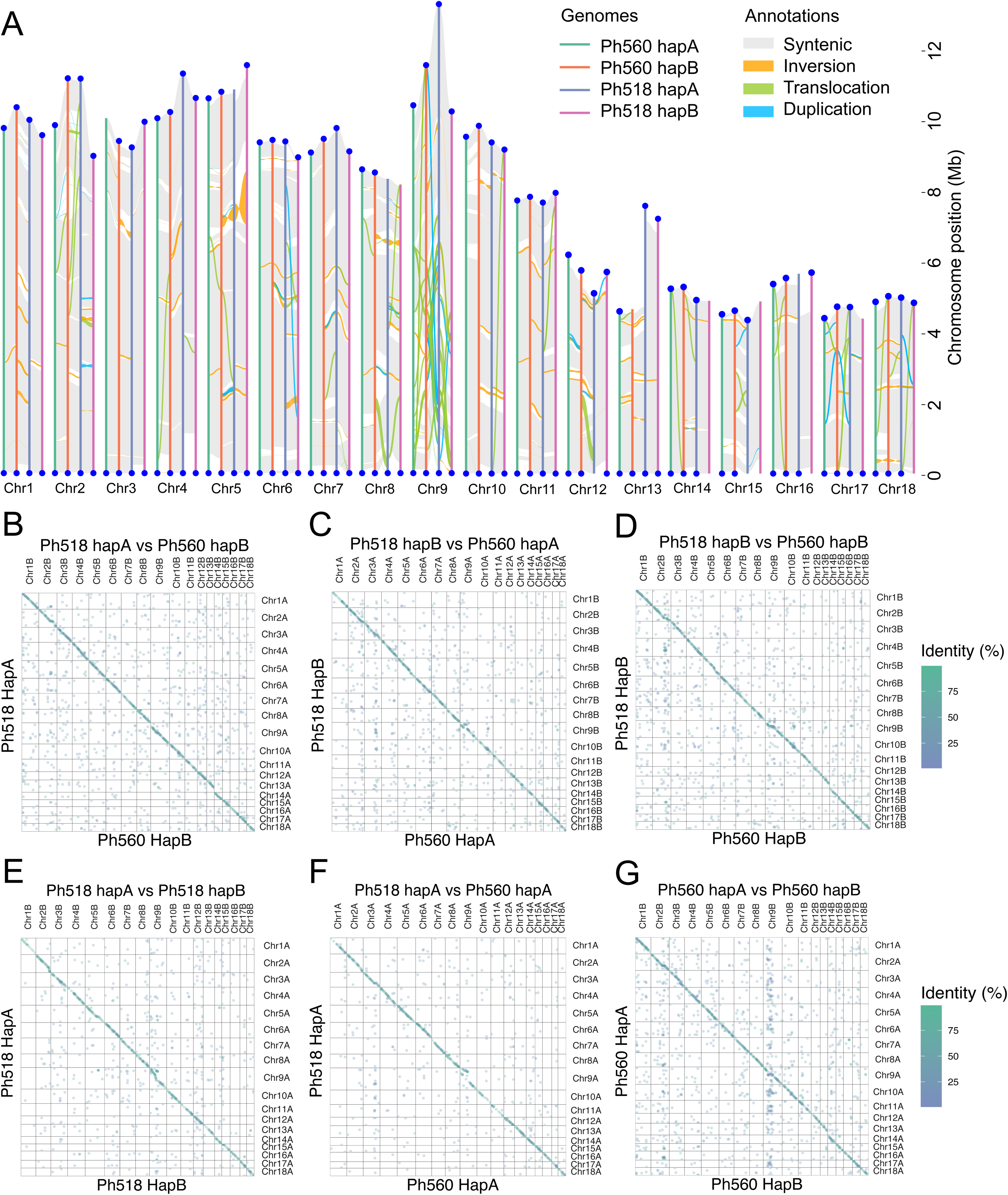
Pairwise whole-genome alignments of haplotype-resolved *Puccinia* isolates Ph560 and Ph518. (A) Chromosome-scale synteny plots between haplotypes A and B of Ph560 (left) and Ph518 (right). Each vertical pair of bars represents the full set of chromosomes for hapA (left) and hapB (right), with colored ribbons connecting syntenic regions. Syntenic regions are shown in grey. Ribbon orientation reflects alignment direction (co-linear or inverted), and crossing patterns or overlap indicate structural rearrangements such as inversions, duplications, or translocations. Identified telomeres are highlighted as blue dots at chromosomal distal ends. (B-G) Pairwise alignments of whole haplotype assemblies using minimap2 (v2.26, asm5 preset). All pairwise combinations of haplotypes from Ph560 and Ph518 were aligned at the chromosome scale. The x-and y-axes correspond to the genome coordinates of the two haplotypes being compared. Diagonal lines represent co-linear alignments; off-diagonal or repeated blocks indicate structural rearrangements or duplications. Color shading denotes sequence similarity.

**Table 1:** Assembly metrics of the two *Puccinia hordei* isolates Ph518 and Ph560

| Parameters | Ph518A | Ph518B | Ph560A | Ph560B |
| --- | --- | --- | --- | --- |
| Chromosomal scaffolds | 18 | 18 | 18 | 18 |
| Chromosomal length (bp) | 147501061 | 141623950 | 142632809 | 143952256 |
| Longest chromosome (bp) | 13261992 | 11537974 | 10599434 | 11537491 |
| GC content (%) | 41.6 | 41.6 | 41.6 | 41.6 |
| Repeat content (%) | 72.2 | 72.7 | 71.8 | 72.9 |
| Complete BUSCOs (%) | 93.1 | 91.5 | 95.0 | 94.3 |
| Single-copy BUSCOs (%) | 85.3 | 86.2 | 91.5 | 90.4 |
| Duplicated BUSCOs (%) | 7.8 | 5.3 | 3.5 | 3.9 |
| Fragmented BUSCOs (%) | 0.9 | 1.0 | 0.9 | 0.8 |
| Missing BUSCOs (%) | 6.0 | 7.5 | 4.1 | 4.9 |

A high coverage (130x) of PacBio HiFi reads with the hifiasm HiFi-only mode was used to generate initial haplotype assemblies for Ph518. After deduplication, contigs from the two haplotypes contained a total of 61 telomeres at contig ends, indicating highly-continuous assembly of these contigs that approached chromosomal level. Contigs were then scaffolded by using the Ph560 Hi-C reads followed by homology-based genome correction, which yielded 18 chromosomes for each haplotype (Figure 2A). BUSCO completeness analysis revealed slightly reduced quality compared to the Hi-C scaffolded Ph560 assemblies, with 93.1% completeness for Ph518A and 91.5% for Ph518B. The combined Ph518 haplotypes achieved 95.1% complete BUSCOs (Table 1). Genome alignment with Pt15 haplotype1 reference revealed clear chromosome-to-chromosome collinearity for all four *P. hordei* haplotypes (Supplementary Figure 2).

### LTR aging and gene structure annotation

The four haplotype assemblies were soft-masked using RepeatMasker with customized repeat libraries. All haplotypes were highly repetitive, with over 70% of each genome composed of repetitive elements (Supplementary Figure 3A). Among these, LTR retrotransposons were the most abundant, accounting for over 38% of each genome. To investigate whether these repeats accumulated gradually or underwent a sudden expansion during the evolutionary history of *P. hordei*, LTR repeat alignments were extracted from the output of RepeatMasker and then analyzed. The results indicated that all four haplotypes experienced an LTR-retrotransposon burst approximately 3 million years ago (Supplementary Figure 3B).

Gene prediction based on both transcriptome and orthologous protein evidence annotated over 11,000 genes per haplotype (Table 2). Subsequent secretome prediction identified approximately 1,500 putative secreted proteins in each haplotype (Supplementary Tables 2A-D), among which around 800 candidate effectors were further predicted.

**Table 2:** Summarized results of gene structure annotation, and predictions of secreted proteins and effectors for the four *Puccinia hordei* haplotypes.

|  | Ph518A | Ph518B | Ph560A | Ph560B |
| --- | --- | --- | --- | --- |
| **Gene structure annotation** |  |  |  |  |
| Number of genes | 11493 | 11485 | 11194 | 11065 |
| Mean gene length (bp) | 1729 | 1751 | 1779 | 1733 |
| Genome covered by genes (%) | 13.5 | 14.2 | 14.0 | 13.3 |
| **Secretome prediction** |  |  |  |  |
| Number of secreted proteins | 1532 | 1547 | 1511 | 1457 |
| Percentage of total proteins (%) | 13.3 | 13.5 | 13.5 | 13.2 |
| **Effector prediction** |  |  |  |  |
| Candidate effectors | 881 | 886 | 859 | 809 |
| Percentage of secreted proteins (%) | 57.5 | 57.3 | 56.8 | 55.5 |

### Comparative genomics reveals synteny and structural diversity between haplotypes

To investigate genome-wide similarities across different haplotypes, we performed syntenic and pairwise comparisons among the four haplotype assemblies (Figures 2A-G). We observed strong collinearity patterns, reflecting high structural conservation between haplotypes (Figure 2A). Most chromosome pairs exhibited near end-to-end alignments and syntenic blocks, indicating successful haplotype resolution and consistent chromosomal scaffolding, particularly for Ph518. To compare haplotypes in terms of sequence similarity, a heatmap was generated based on the percentage of alignments with sequence identities greater than 50%. The results revealed that Ph518A and Ph518B were the most conserved pair, followed by Ph518A and Ph560A, while Ph518B and Ph560B were the most divergent (Supplementary Figure 4 and Supplementary Table 3). Despite the overall high level of structural conservation, clear instances of haplotype-specific and isolate-specific structural variations (SVs) could also be observed (Figure 2A).

Within Ph560 (Figure 2A), while some chromosomes such as Chr1, Chr3, Chr7, and Chr14 maintained strong haplotype co-linearity, several others exhibited prominent haplotype-specific variations. Chr2 contained a large syntenic gap along with partial duplications between haplotypes. Chromosomes 5, 8, 12, 13, and 18 displayed more modest rearrangements, including partial inversions or translocations. In Ph518, consistent collinear patterns were found for a majority of the homologous chromosomes, whereas large inversions, translocations and duplications were present in Chromosomes 5 and 8. While many chromosomes retained consistent synteny between Ph560 and Ph518 (e.g. Chromosomes 11, 14 and 16), some showed strong isolate-specific rearrangements or differing haplotype architectures (e.g. chromosomes 1, 3, 12,15, and 17). Inter-isolate comparisons between Ph560 and Ph518 highlighted a mixture of shared and lineage-specific structural divergence, consistent with their highly divergent virulence profiles.

The most dramatic difference and SVs were observed for Chr9 in both inter-and intra-isolate syntenic comparisons, where large, complex structural rearrangements disrupted central synteny across all haplotypes, suggesting a major haplotype-specific inversion or breakpoint spanning a substantial portion of the chromosome. We observed greater haplotype asymmetry in Ph518 with the presence of abundant translocations, fragmented synteny across its central region in Ph560, and extensive duplications between Ph560 and Ph518 (Figure 2A). Genome alignment also indicated that Chr9 consistently showed the highest number of duplicated alignment blocks and misaligned segments, even in the most similar haplotype pairs (Figures 2E-F). These patterns were not seen on other chromosomes to the same extent, suggesting that Chr9 has undergone the most dramatic structural divergence both within and between isolates. It is possible that Chr9 is a genomic hotspot of plasticity, potentially harboring lineage-specific expansions or gene family diversification relevant to adaptive processes in *P. hordei*.

Orthogroup clustering analysis across six *Puccinia* species demonstrated a high level of conservation. Of the total 11,194 proteins in *P. hordei*, 93.1% (10,422) were assigned to orthogroups, while only 6.9% were classified as singletons, indicating highly conserved protein families with other *Puccinia* species (Supplementary Table 4). Of the 7,426 orthogroups identified in *P. hordei*, 258 were species-specific, potentially representing unique genomic adaptations related to host specialization or pathogenicity. GO enrichment of these *P. hordei*-specific orthogroups revealed significant enrichment in functional categories such as “rRNA processing” and “protein transport”, along with “vesicle-mediated transport”, “signal transduction”, and “ubiquitin-dependent catabolism”, suggesting roles in intracellular trafficking and signaling pathways (Supplementary Figure 5A). Phylogenetic analysis of the six *Puccinia* species using single-copy orthogroup proteins was used to trace their evolutionary history. Interestingly, the wheat leaf rust pathogen *Pt* is more closely related to *P. hordei* than to other *Puccinia* species infecting wheat, despite distinct host specializations between *Pt* and *P. hordei* (Supplementary Figure 5B). Collinearity analysis among the four *P. hordei* haplotypes was conducted using orthogroup clustering results. Consistent with the genome-wide syntenic patterns (Figure 2A), extensive synteny was observed across the haplotypes, with conserved blocks spanning most of each genome, suggesting a shared genomic architecture (Supplementary Figure 6). Disrupted or shuffled regions reflected small structural variations between haplotypes, where Chr9 again showed strong evidence of duplications in orthogroup genes (Supplementary Figure 6).

### Chromosome 9 exhibits unique repeat structure, compartmentalization, and genomic composition

Prompted by the observation of haplotype-specific structural variations, we investigated whether Chr9 exhibits distinct genomic and architectural features that set it apart from the rest of the *P. hordei* genome. We hypothesized that Chr9 might have undergone unique evolutionary processes, including repeat-mediated expansion, reorganization, and functional domain segregation. To test this, we conducted comparative analyses across genome structure, Hi-C compartments, and genomic content, focusing on both whole-genome and Chr9-specific patterns.

We first assessed intra-chromosomal architecture for Chr9 in Ph560 haplotypes using a self-alignment plot (Figure 3A and Supplementary Figure 7C). Chr9A displayed dense and regular diagonal signals, with tandem and inverted repeat expansions. These repeat arrays were widespread and prominent across Chr9A, shown as relatively sparse self-similarity patterns consistent with other genome/chromosome regions (Figure 3A and Supplementary Figure 8). However, a notable and intense self-similarity region of Chr9 was observed spanning ∼3Mb adjacent to the putative centromere, which is enriched with simple repeats (Figure 3A, Supplementary Figure 7C and Supplementary Figure 8). In the Hi-C maps (Figure 3B and Supplementary Figure 7D), the same regions show elevated local interaction frequency, consistent with physical compaction and structural insulation. This repeat block corresponds precisely to a sharp E1 eigenvector transition (Figure 3B and Supplementary Figure 7D), suggesting they acted as compartment boundaries. Gene density is markedly reduced within this block, yet candidate effectors tend to cluster near the flanks and centromere expanded region (Figure 3A and Supplementary Figure 7C). The localization and sequence content of this region suggest it may serve as a repeat-mediated architectural module, contributing to domain insulation and possibly shaping the chromatin environment around the centromere. The simple repeat compositions may also promote structural variation and serve as a genome organization anchor.

**Figure 3:**
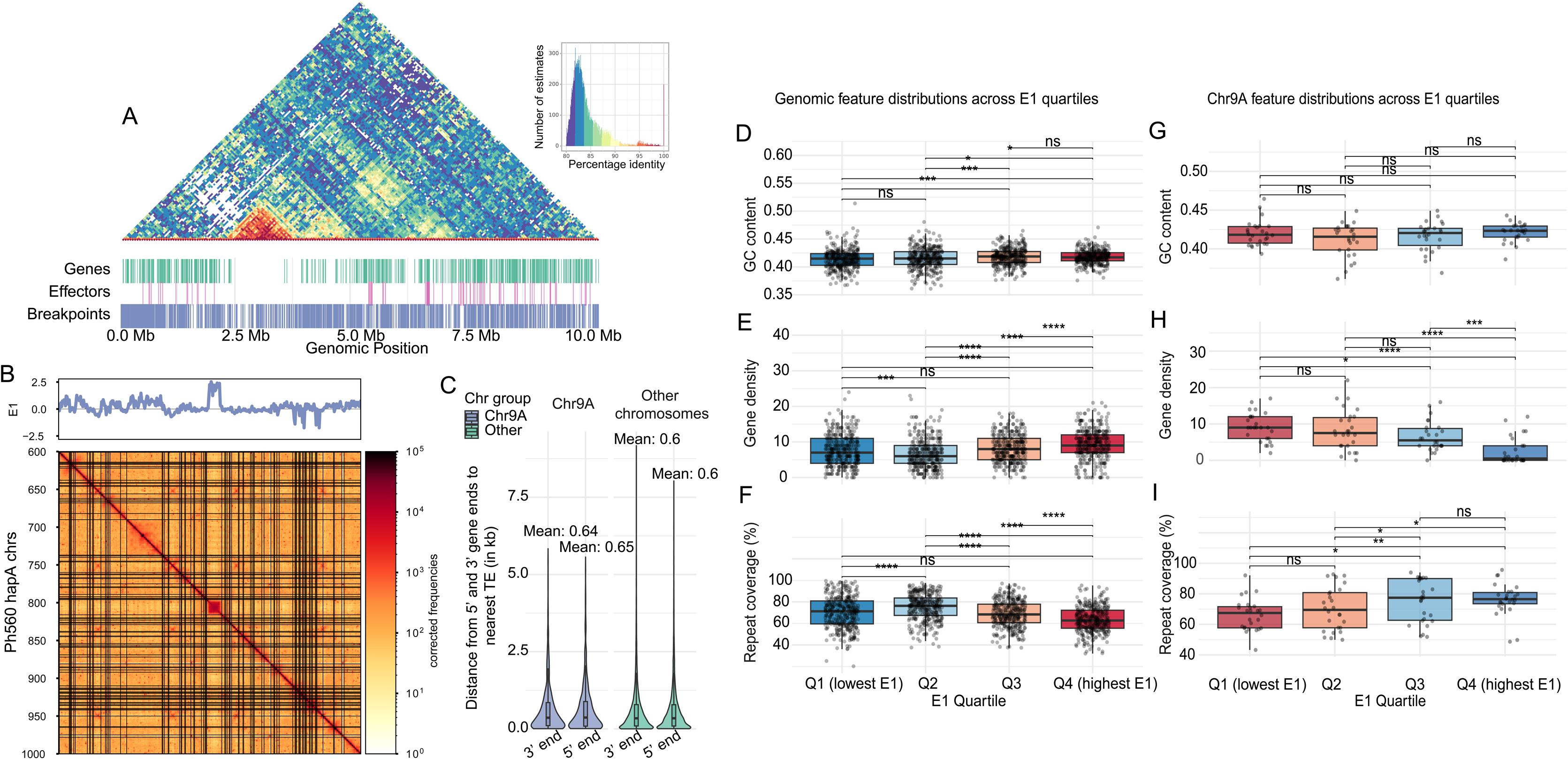
Structural, spatial, and compositional genome features of Ph560 Chr9A. (A) Whole-chromosome self-alignment of Chr9A reveals widespread tandem and inverted repeat arrays. Gene and predicted effector annotations (green and purple ticks) and inter-isolate alignment between Ph560 Chr9A and Ph518 Chr9A are shown along the axes. SV coordinates between Ph560 Chr9A and Ph518 Chr9A were represented by breakpoints shown as gaps. The upper display indicates percentage identity distribution of alignments. (B) Hi-C contact matrix and first eigenvector (E1) of Chr9A in Ph560, highlighting pronounced compartment-like domains spanning bins 600-1000 (chromosomes 7-10). Heatmap indicates corrected contact frequencies. Crosses reflect centromere locations. (C) Distances from 5’ and 3’ gene ends to the nearest TE, comparing Chr9A to all other chromosomes. Chr9A genes are slightly more distant from TEs on average. (D-I) Genomic features in compartments of the Ph560 hapA genome. (D-F) Genome-wide distributions of GC content, gene density (number of genes), and TE coverage across Hi-C E1 quartiles. Q1 (lowest E1) is associated with low GC/gene content and high TE load, while Q4 (highest E1) shows the opposite. (G-I) Compartmental distributions of the same genomic features specifically for Chr9A. In contrast to the genome-wide trend, Chr9A exhibits an inverse relationship where gene density increases, and TE coverage decreases from Q1 to Q4. 100kb-window was used. Pairwise t-tests were conducted (ns not significant; * *p* < 0.05; ** *p* < 0.01; *** *p* < 0.001; **** *p* < 0.0001).

We aligned haplotype A or B genomes of Ph560 and Ph518 to assess haplotype-specific structural divergence for Chr9. Extensive alignment breakpoints (2.5 kb window) occurred between the two isolates, at or near these repetitive regions (Figure 3A and Supplementary Figure 7C). Gene annotation overlays suggested that candidate effector-like genes tended to cluster around alignment breakpoints in Chr9. These hotspots of SV breakpoints may drive effector diversification, leading to virulence evolution in *P. hordei* given the highly distinct virulence patterns of the two isolates.

We next explored higher-order chromatin organization using Hi-C eigenvector decomposition (E1 values). In the Ph560 contact matrix, the genome exhibited well-defined alternating E1 domains including Chr9A, which approximately spanned bins 650 to 850 (Figure 3B and Supplementary Figure 7D), suggesting a clear bipartite structure and compartmentalization. To examine whether compartmentalization might affect the spatial relationship between genes and nearby TEs, we examined the distances (both 5’ and 3’ ends) of genes to the nearest TE and compared Chr9A to all other chromosomes. On average, Chr9A genes had slightly longer TE distances (∼0.63-0.64 kb) than genes on other chromosomes (∼0.61 kb) (Figure 3C). Despite lacking statistical significance, the TE-gene spacing difference can be considered dramatic and a strong distinguishing feature of Chr9A because of the above-70% TE/repetitive genome (Supplementary Figure 3A) and the large span of a simple repeat enriched region affecting gene spacing.

To dissect how Ph560 genome compartmentalization relates to compositional features, we examined GC content, gene density, and TE coverage across E1 eigenvector quartiles (Q1-4). When genome features were aggregated at the chromosome scale, Chr9A and Chr9B consistently ranked among the lowest in gene density (∼65 and 63 genes/100kb, respectively) and highest in TE coverage (∼72% and ∼77.7%, respectively) compared to all other chromosomes (Supplementary Figures 7A-B). Interestingly, the relationship between chromatin compartment and genomic features on Chr9A is the opposite of what was seen for the rest of the genome. In other chromosomes, high E1 compartments (Q4) are associated with elevated GC content, gene density and reduced TE content (Figures 3D-F). However, in Chr9A this pattern is reversed - GC remained similar across quantiles (Figure 3G), Q3-Q4 bins exhibited significantly reduced gene density and increased TE coverage (Figures 3H-I), while lower E1 bins (Q1-Q2) showed relatively higher gene density and lower TE content (Figures 3H-I). This pattern suggests a clear spatial differentiation between gene-rich, GC-rich, TE-poor domains and gene-poor, GC-poor, TE-dense regions. The polarity inversion highlights a distinctive chromatin-sequence relationship on Chr9A, suggesting that its compartmental architecture may serve as an alternate regulatory or structural role, potentially shaped by extensive repeat integration and genome rearrangement.

On the contrary, the quartile-dependent distribution of features was not clearly observed in Chr9B (Supplementary Figures 7E-J). The observation that Chr9B exhibits E1-compartmental trends that differ from those of Chr9A suggests allelic divergence in higher-order chromatin structure and associated genomic features. In Chr9A, high E1 (Q4) bins are gene-rich and TE-poor, while low E1 (Q1) bins are TE-rich and gene-poor. However, this polarity is flipped or weakened in Chr9B, where Q1 bins remain TE-rich but Q4 bins no longer display a strong gene enrichment and instead maintain relatively high TE content. This discordance may reflect structural polymorphisms or epigenetic variability that decouple E1 signal from functional content in one haplotype. Such intra-individual haplotype-specific chromatin divergence has implications for differential gene regulation, stability of chromatin domains, and potential adaptation through structural variation.

### Population genomics reflects reproductive structure and lineage divergence of a continental *P. hordei* population

No study has clearly delineated clonally derived lineages from those shaped by sexual recombination within a *P. hordei* population. We selected 41 *P. hordei* isolates from a continental collection of pathotypes representing those detected in annual pathogenicity surveys in Australia across a time span of 54 years (Cotterill et al., 1995; Park, 2003). With the exception of one isolate that was recovered from the alternate host *O. umbellatum* (Ph577), all were recovered from leaf rusted barley. Three of the isolates recovered from infected barley were discovered as spontaneous virulence gain mutants in greenhouse testing, while the remaining 37 were sourced from leaf rust infected barley crops (Supplementary Table 1).

To investigate the underlying population structure and infer potential reproductive histories of *P. hordei*, we conducted genome-wide SNP analyses using reference assemblies from both haplotypes of Ph560 to assess the consistency of clustering, co-ancestry, and introgression patterns. Principal component analysis (PCA) revealed similar major axes of variation regardless of haplotype reference genome. In both cases, PC1 and PC2 captured clear separation between major genetic groups, while individuals including Ph518, Ph577 and Ph640 appeared haplotypically divergent (Figures 4A-B; Supplementary Table 1). To interpretate cluster composition, we applied k-means clustering (k = 5) to PC1-PC2 coordinates, which identified five genetically distinct groups for both haplotypes (Figure 4A; Supplementary Table 1). While these clusters are algorithmically derived and do not reflect predefined populations, they correspond broadly to discrete genetic groups also identified in ADMIXTURE (Figure 4C), co-ancestry matrices (Supplementary Figure 9), phylogenetic relationships (Figures 4D-E) and SplitsTrees (Supplementary Figure 10). The isolate Ph518 appeared phylogenetically distinct in PC3-PC4 in both haplotype references, supporting its role as an outgroup consistent with its broad avirulence for most *Rph* genes.

**Figure 4:**
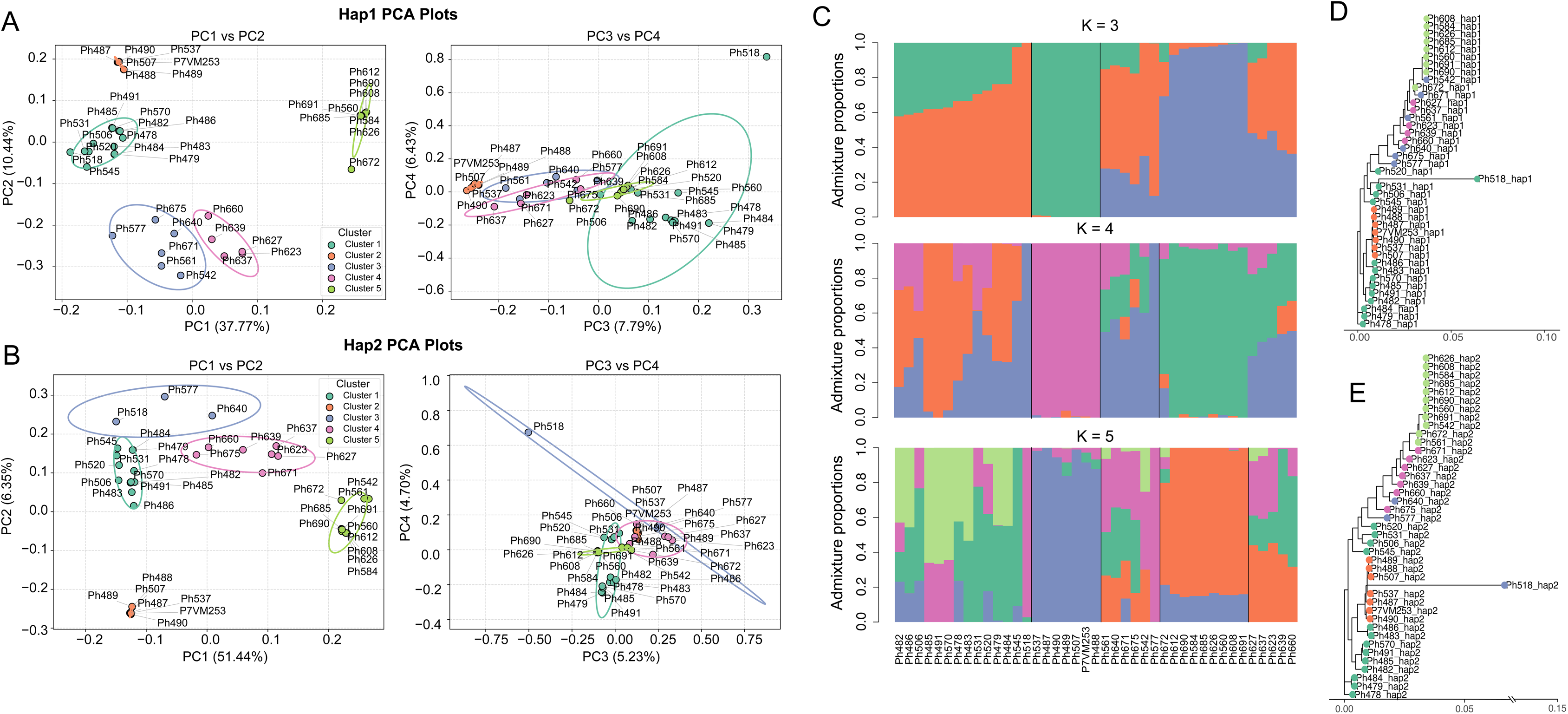
Population structure analyses using SNPs called against both haplotype A and haplotype B reference genomes to assess the consistency of clustering, co-ancestry, and introgression patterns. (A and B) Principal component analysis (PCA) revealed similar major axes of variation regardless of reference genome. K-means clustering (k = 5) to PC1 and PC2, which grouped individuals into five clusters. (C) ADMIXTURE analysis based on haplotype A at K = 3-5 recovered the same five population clusters across both haplotypes, showing high consistency in cluster assignment for clonal groups and revealing mixed ancestry in intermediate individuals. (D and E) Phylogenetic inference of *P. hordei* isolates.

Several isolates (e.g., Ph487, Ph488, Ph489, Ph490, Ph507 and P7VM253 from Group 2 of both haplotypes; Supplementary Table 1) grouped tightly in PCA space (Figures 4A-B) and showed nearly identical ADMIXTURE ancestry profiles at K = 3-5 (Figure 4C), indicating that they likely represent a clonally propagated lineage. Isolates from this lineage also formed a compact cluster in the SplitsTree network with minimal reticulation (Supplementary Figures 10A-B) and exhibited extremely high co-ancestry values in both hapA-and hapB-reference based matrices (Supplementary Figures 9B and 9D). A second clonal group was also identified among Ph685, Ph690, Ph691, Ph584, Ph560, Ph608, and Ph612, all of which displayed tight PCA clustering of both haplotypes (Figures 4A-B), ADMIXTURE homogeneity and tree networks (Supplementary Figures 10A-B). These patterns support long-term (over 50 years) asexual and sexual reproduction of two independent progeny groups of *P. hordei*. The two distinct clonal lineages are consistent with long term data on pathotype incidence, where it is likely that isolates of the first group (i.e. Ph488 and Ph487 collected in 1966 and 1969, respectively) were dominant in the 1960s and 1970s (Supplementary Table 1). There was likely another group typified by Ph482 that dominated in the 1980s/1990s (Supplementary Table 1) due to virulence on *Rph4* and cultivation of barley cultivar Grimmett which carries this gene (Cotterill et al., 1995; Park, 2003), and a derivative pathotype typified by isolate Ph491 after 2001 due to its virulence on *Rph12* and cultivation of several barley cultivars carrying this gene (Park, 2003; RF Park unpublished). The minor genetic difference for these old isolates such as Ph482, Ph487 and Ph488 could be revealed by within group admixtures patterns (Figure 4C) and additional PCA levels (Supplementary Figure 11).

In contrast, individuals with intermediate or dispersed positions across PCs falling between genetic groups and being highly-admixed (e.g. Ph623, Ph627, Ph639 and Ph660 from Group 4; Ph672 from Group 5) (Figures 4A, C and D) suggest signatures of historical recombination or hybrid origin. The f-branch statistic (fb), computed independently for each reference haplotype, provided further evidence of introgression. In hapA-based analysis (Supplementary Figure 9A), Group 5 showed elevated fb values from Group 4 (fb = 0.98), with additional contributions from Group 1 (fb∼0.9) and Group 2 (fb∼0.6), indicating multiple historical gene flow events. In the hapB f-branching (Supplementary Figure 9C), analysis indicated significantly stronger gene flow between Group 5 and Group 3 (fb ≈ 0.9), suggesting strong allele sharing between Group 5 with Group 3. The two haplotype patterns further indicate recombination of different donor lineages. Further support for this complex ancestry comes from the co-ancestry heatmap, where Ph660 as a Group 4 isolate exhibited high co-ancestry with the Group 5 clonal block and clustered tightly within it (Supplementary Figure 9B), and Ph518 and Ph640 from Group 3 with high co-ancestry relationships to Group 5 isolates (Supplementary Figure 9D). Therefore, Ph660 and Ph518/Ph640 may be more likely related to origins of parents or introgressed individuals, and similar to the genetic makeup of hapA or hapB Group 5 isolates. Together, these findings strongly support a recombinant origin of Ph isolates from Groups 3-5, involving donor or contributions from multiple groups and highlight the presence of recent or historical recombination among a subset of isolates, in contrast to the clearly clonal structure observed in other groups (Groups 1 and 2). Combing data from surveys and pathogenicity tests, hapA and hapB Group 5 isolates (collected in the early 2000s) (Supplementary Table 1) are considered the original founding group, from which Groups 3 and 4 were derived.

### Reproductive lineages inferred from Chr9 haplotype k-mer and mating-type divergence

Reproduction and mating-type genes are strong determinants of stages of fungal lifestyles and population structures. To further investigate the historical collection of *Ph* isolates, we used mating-type genes from *Pt* and *Pst* as queries to search for homologs in *P. hordei*. The P/R and HD loci were identified to be unlinked, with locations on separate chromosomes. For the P/R locus, sequence blasting identified the *STE3.1* gene, previously speculated to be uninvolved in mating compatibility in other rust fungi (Luo et al., 2024), in all four *P. hordei* haplotypes on Chromosome 1, showing 100% coding sequence identity within isolates and 99.4% identity between isolates. The *STE3.2* and *STE3.3* homolog genes were located on Chromosomes 9A and 9B of Ph560 but haplotypically swapped in Ph518. Despite this, the coding sequences of *STE3.2* and *STE3.3* across haplotypes of Ph560 and Ph518 are identical. This observation is consistent with the extensive inter-/intra-haplotype SVs on Chr9 where clear inversions were found for the *STE3.2* and *STE3.3* loci at ∼4.8 Mb positions (Figure 2A). Orthogroup clustering with other *Puccinia* species correctly grouped these *P. hordei STE3* genes into STE3.1 (Supplementary Figure 12A) and STE3.2/3.3 (Supplementary Figure 12B) clusters. Orthologous *STE3* genes from *Pt* showed the closest evolutionary relationship to *Ph*. In contrast, multiple alleles of the HD locus were identified on Chromosome 4 with different *bW-HD1* and *bE-HD2* alleles present in each haplotype. The *bW-HD1* and *bE-HD2* alleles were closely linked, encoding polypeptides of ∼600 and ∼370 amino acids, respectively. Grouping of orthologous genes also correctly clustered these *P. hordei bW-HD1* (Supplementary Figure 12C) and *bE-HD2* (Supplementary Figure 12D) alleles into two different clusters, with *Pt* still the closest related species.

Because of the distinct variation features and P/R loci of Chr9, we conducted Chr9-specific k-mer analysis and found a clear bipartition pattern among *P. hordei* isolates across both haplotypes (Figures 5A-B). Clustering of the top 100 most variable k-mers independently extracted from Chr9A and Chr9B divided the isolates into two primary groups (Figures 5A-B). These groups were non-overlapping between the two chromosomes where isolates with high k-mer diversity on one haplotype showed low signal on the other. When similar k-mers were collapsed into clusters (Figures 5C-D), the groupings showed dramatic partitioning, with one most similar k-mer group for each of the haplotypes (49 k-mers and 88 k-mers, respectively; Supplementary Table 5). This reciprocal haplotype-specific pattern suggests inheritance of structurally divergent Chr9A and Chr9B haplotypes across the population, likely maintained by recombination suppression or structural incompatibility. Given the dikaryotic nature of *P. hordei*, these results indicate that only one haplotype is highly represented per Chromosome 9 in a given isolate, and that the divergence is sufficient to stratify the population.

**Figure 5:**
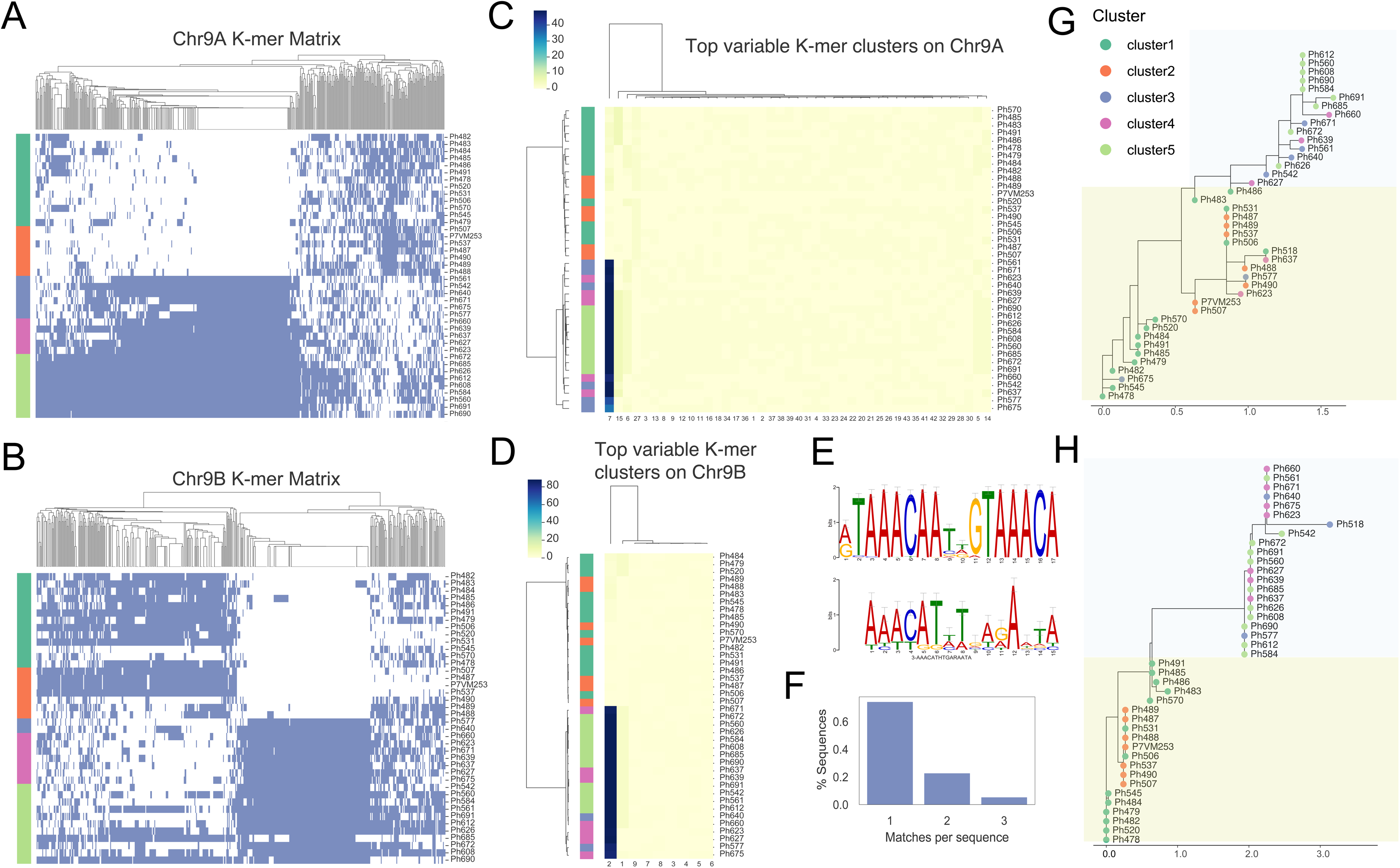
Chr9 k-mer signals reflecting mating-type linked genomic divergence that separates lineages. k-mer clustering revealed population structures that could map onto known mating-type gene divergence consistent with reproductive mode of *P. hordei*. (A and B) Distribution of sample-wise k-mers present across Chr9A (A) and Chr9B (B). (C and D) The most variable (top 100) k-mer features on each haplotype. To interpret the genomic content underlying these k-mer differences, the most variable k-mers were clustered by sequence similarity resulting in a reduced representation matrix. Collapsed k-mer clusters recapitulate the same two-group structure seen in the full matrix and identify specific clusters with highly skewed distributions across isolate groups. Color key showing k-mer sequence numbers. (E) Clustered k-mer sequences are enriched in a motif related to forkhead/winged-helix (FKH) motifs. (F) Motif matches per sequence. Most k-mer clusters show a single dominant motif match per sequence. (G and H) Phylogenetic Trees of *STE3.2* and *STE3.3* coding sequences. *STE3.2* on Chr9A (G) and *STE3.3* on Chr9B (H) both yielded trees with two dominant clades (grey and yellow shaded) consistent with the k-mer clusters. Colors represent isolates from 5 major genetic groups assigned forementioned.

Among these two groups, the first is composed of older isolates exclusively belonging to Groups 1 and 2 (Figures 5A-B; Supplementary Table 1). These groups also exhibit highly compact clustering in PCA and co-ancestry matrices (Figure 4, and Supplementary Figures 9B and 9D), consistent with clonality. These isolates showed higher homogeneous Chr9A k-mer patterns (Figures 5A and 5C), minimal intra-group variation and collapsed signal in the alternative haplotype (Figures 5B and 5D). Phylogenies of *STE3.2* and *STE3.3* (Figures 5G-H) supported this grouping, in which these isolates exhibited limited allelic diversity in one of the mating-type loci, with some loci nearly invariant. Hence, this first group is best interpreted as asexually maintained, representing long-term clonal lineages with retained haplotype-specific content for either Chr9A or Chr9B, but not both. Notably, some isolates (i.e., Groups 3 and 4; Figures 5C and 5G) appeared mixed or intermediate. However, they still formed a consistent clade (Figure 5G), possibly reflecting recombinant haplotypes or recent gene conversion, consistent with rust mating-type biology (Holden et al., 2023).

In contrast, the second group included isolates from Groups 3-5, which displayed substantial k-mer diversity on both Chr9A and Chr9B. This group exhibited more balanced and variable signals across both chromosomes, consistent with recombined haplotype content. Phylogenies of *STE3.2* and *STE3.3* coding sequences showed higher diversity and the likely maintenance of divergent mating-type alleles (Figures 5G-H). Additionally, the presence of both alleles in separate clades suggests that recombination between haplotypes has been retained. These isolates also showed clear gene flow signatures with Group 5 likely as the founder in f-branch and co-ancestry analysis (Supplementary Figure 9; Supplementary Table 1). Together, these data support that Groups 3-4 are sexually derived, with Chr9 diversity shaped by mating-type allele divergence, recombination, and haplotype mixing.

*De novo* motif discovery among the most variant k-mers from both Chr9A and Chr9B recovered a conserved motif resembling forkhead/winged-helix DNA-binding sequences (Figure 5E), often associated with transcriptional regulators such as *Fkh1* in *Saccharomyces cerevisiae*. This suggests a possible haplotype specific function in regulating mating-type linked processes in the sexually derived isolates. Most k-mers with significant matches carried only one motif hit (Figure 5F), suggesting these are discrete, non-repetitive sequences, possibly located within regulatory or structural domains.

### Genome-wide CNV profiling reveals clonal divergence and progressive shift of lineages

To uncover genomic structural variants underpinning phenotypic divergence, we looked further at sequence divergence across the 41 *P. hordei* isolates. Prompted by the structural variation signatures detected in the whole-genome comparisons (Figure 2), especially the dominant duplications observed in Chr9, we specifically profiled genome-wide CNVs across 40 *P. hordei* isolates (long-reads based Ph518 excluded) representing the pathotypes sampled over a 54-year period (Figure 6A).

**Figure 6:**
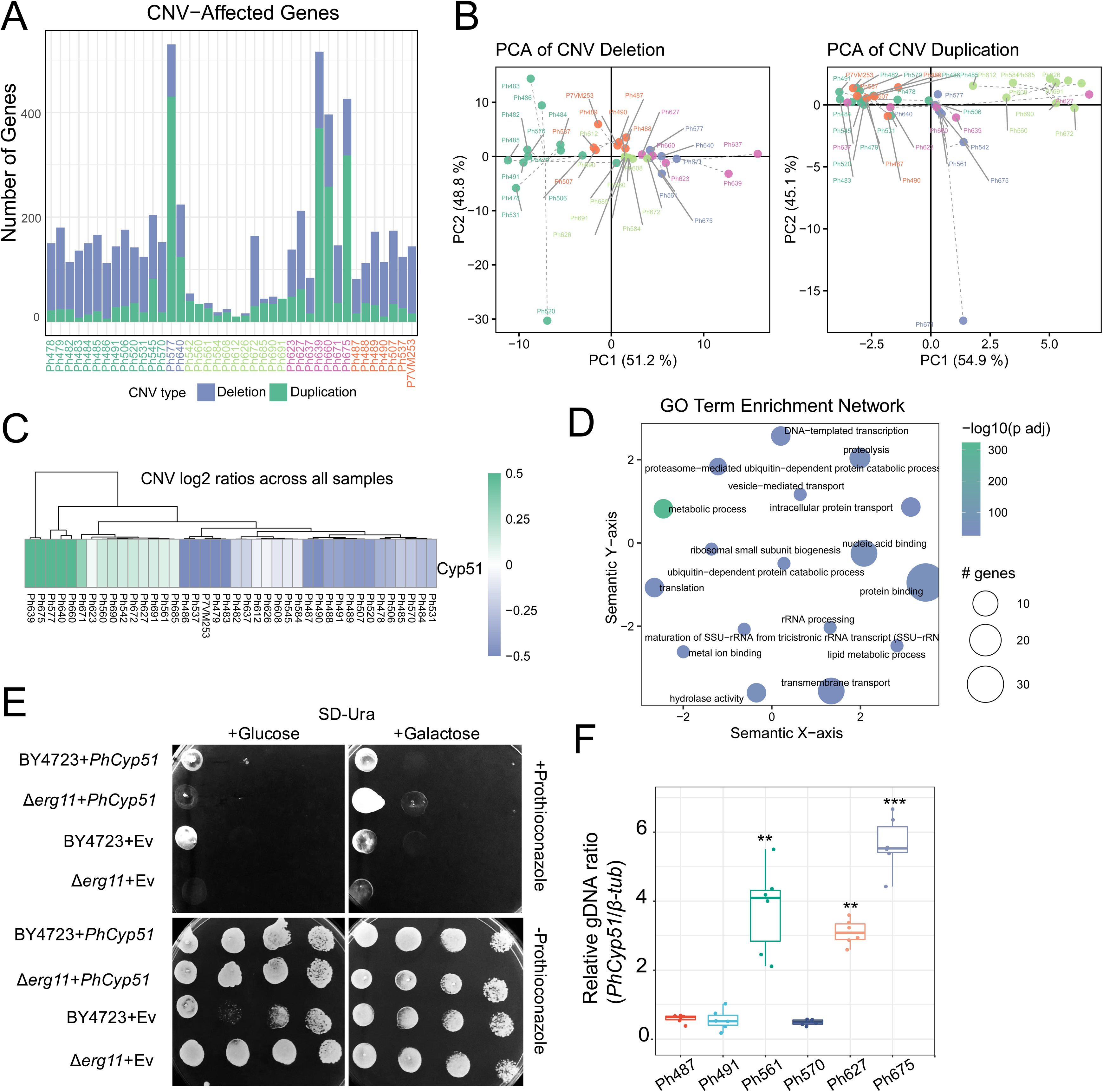
Genome-wide CNV landscape highlights structural divergence across *P. hordei* isolates and links *Cyp51* to fungicide insensitivity. (A) Total number of genes affected by deletions and duplications across *P. hordei* isolates, revealing widespread CNV activity across lineages. Colors indicate previous assigned genetic groups of *P. hordei* isolates. (B-C) Principal component analysis (PCA) of CNV profiles shows a clear separation between older and recently emerged isolates based on deletions (B) and duplications (C), indicating a clear shift toward the increasing of CNV duplication type in newly derived lineages (clear shift of variation types observed between old and new isolates collected across 54 years) (D) Heatmap showing CNV directly affects *Cyp51* across isolates more recently emerged. CNV values were log 2 confirmed. (E) GO enrichment analysis of duplicated genes shows significant overrepresentation of terms related to metabolism, intracellular transport, protein complex organization, and transcriptional regulation. Circle size represents the number of genes per term; color intensity indicates significance. (F) Functional validation of *PhCyp51* in a yeast *erg11* deletion strain confirming its ability to confer resistance to the DMI fungicide prothioconazole. BY4723 WT background and empty vector were used as control. (G) Quantitative PCR showing elevated *Cyp51* gene CNV status in selected isolates. gDNA fold-change of *PhCyp51* relative to *beta-tubulin*. Pairwise T-test with ** *p* < 0.01; \*\*\**p*< 0.005. Showing is representative of two independent gDNA extractions showing the same results.

PCA of gene-CNV level (log CNV ratios) revealed a striking temporal trajectory (Figures 6B-C). Isolates detected early in the survey period (e.g., Ph487-490; Supplementary Table 1) were characterized by widespread deletions and low CNV levels, while more recent isolates (e.g., Ph660, Ph672, Ph675; Supplementary Table 1) displayed substantial CNV duplications across multiple chromosomes. This shift reflects ongoing clonal diversification, with genome remodeling accumulating in more recent genotypes.

To evaluate *P. hordei* chromosomes and isolates most impacted by CNV, we computed average CNV intensity (log ratios) per chromosome in 10 kb bins across all isolates for both haplotypes. The heatmaps (Supplementary Figures 13A-B) revealed clear lineage-specific CNV signatures that were consistent within the previously assigned groups (Supplementary Figure 9). Isolates from Group 5 exhibited clear duplications across a majority of chromosomes, especially in Chr9 of both haplotypes, suggesting high-copy duplications specific to this group. These patterns indicate that the accumulation of CNVs, particularly duplications, is a hallmark of recent clonal diversification, with Group 5 lineages undergoing the most extensive genome remodeling. An exception was Ph488, which although not classified in Group 5, showed high CNV levels in about half of its genome. However, Chr9 in this isolate had clearly not gained CNVs. Because Chr9 is hypothesized as being a hotspot of evolution in *P. hordei* (Figure 3 and Supplementary Figure 7), CNV status still may differentiate Ph488 from Group 5.

TEs have been recently shown as a major genomic driver for CNV in fungal plant pathogens (Stalder et al., 2023; Tralamazza et al., 2024). To explore the genomic drivers of CNV formation, we examined the distribution and distance of TEs relative to CNV loci. We found that CNV loss (deletions) was significantly closer to TEs than duplications and normal loci (Supplementary Figure 13C). Specific TE classes, including LINE/J-jockey, LTR/Copia elements, low-complexity, simple repeats and unknown categories, showed the strongest spatial association (shorter distances) with deletions (Supplementary Figures 13F, H-I, and M-N) or duplications (Supplementary Figures 13M-N), suggesting that local repeat architecture may facilitate tandem duplications or segmental amplification.

To investigate the functional impact of gene duplications, we performed GO enrichment analysis on genes overlapping high-confidence CNV gains. Broad biological processes such as “metabolic process”, “cellular process”, and “intracellular anatomical structure” were among the most significantly enriched terms (adjusted p < 5×10[³²[) (Figure 6D). More specific categories included “lipid metabolic process”, “intracellular protein transport”, and “protein-containing complex organization”, suggesting duplications affect membrane trafficking and metabolic regulation. Additional enrichment in “carbohydrate transport” and “protein targeting” points to selective expansion of genes involved in nutrient acquisition and cellular remodeling. These results indicate CNV-driven amplification preferentially targets functional modules likely contributing to pathogen adaptation under selection pressures associated with agricultural practices.

### CNV-driven divergence in recent *P. hordei* lineages is associated with fungicide insensitivity

During investigation of CNV-driven genomic plasticity, we identified *Cyp51* as a key gene recurrently duplicated in a subset of isolates (Figure 6C). These isolates were exclusively newly emerged isolates including those belonging to Groups 5 and 4 (Figure 4 and Figures 6A-C). *Cyp51* encodes sterol 14α-demethylase, the primary target of DMI/triazole fungicides. *Cyp51* has only been referred as a candidate for the wheat stripe rust pathogen based on population genome-wide sequence variations, however, neither fungicide insensitivity nor functional assay could confirm this either *in-vitro* or *in-planta* (Cook et al., 2021). In Ph560, there are five identical copies of *Cyp51* (one on Chr9A, and five on Chr9B Supplementary Figure 14), all located in regions that were previously identified as a CNV hotspot in Chr9 of the Ph560 genome. In contrast, only two copies were found in Ph518, showing no variation in protein coding sequences to that of Ph560 (Supplementary Figure 14). Hierarchical clustering of CNV levels (log ratios) across isolates confirmed lineage-specific duplications of *Cyp51* (Figure 6C). In contrast, the gene showed neutral or deletions in earlier diverging isolates (Figures 6A-C). This gene showed consistent copy number gains in isolates with elevated genome-wide CNV levels (Figure 6A). More consistently, these were the same isolates that are considered to be part of the clonal lineage typified by pathotype 5453 P-, of which Ph560 (Supplementary Table 1) is the founding isolate. *In planta* studies under controlled conditions have shown that these isolates are insensitive to several DMI fungicides (Park et al., 2022).

The founding isolate of the 5453 P-clonal lineage, Ph560, was first detected in Western Australia in 2001. It was first detected in eastern Australia the following year, and by 2006 it or mutational derivatives from it had been detected in all barley-growing regions in Australia (Park RF, unpublished), and all of those tested under controlled conditions so far have displayed insensitivity to several DMI fungicides (Chhetri et al., unpublished). To confirm the CNV duplication patterns, we quantified *Cyp51* copy number by qPCR using gDNA of three representative isolates belonging to the 5453 P-lineage (Ph561, Ph627, and Ph675) and three other isolates (Ph457, Ph491, Ph570). As shown in Figure 6F, we observed significantly higher genomic *Cyp51* copies in the three randomly selected 5453 P-lineage isolates in contrast to low copy states in the other three isolates. To test functional relevance, we cloned the *PhCyp51* coding sequence into a galactose-inducible expression vector and introduced it into the *S. cerevisiae* Δ *erg11* mutant strain. In the absence of native ergosterol biosynthesis, *PhCyp51* expression restored growth on galactose media and conferred resistance to the DMI fungicide prothioconazole, confirming functional compatibility and azole insensitivity in a heterologous system (Figure 6E). No growth rescue was observed for the empty vector controls or when grown on non-inducing glucose medium.

## Discussion

This study presents the first haplotype-resolved genome assemblies for *P. hordei*, the causal agent of barley leaf rust, providing a comprehensive structural and functional map of its diploid genome. Using long-read sequencing and Hi-C scaffolding, we reconstructed chromosome-level assemblies for two pathogenically diverse isolates, revealing extensive structural conservation interspersed with haplotype-specific and lineage-specific variation. Recently developed tools were employed to enhance assembly and annotation quality. Hi-C scaffolding was performed using YaHS, which constructs contact matrices with improved contig-joining accuracy and outperforms or matches other scaffolders such as 3d-DNA, SALSA2, and ALLHiC in benchmark comparisons (Hou et al., 2023; Obinu et al., 2024). Gene prediction was conducted using BRAKER3, which integrates GeneMark-ETP and AUGUSTUS with transcriptome and orthologous protein evidence to generate high-confidence gene models (Gabriel et al., 2023). RNA-seq data for annotation were obtained from *P. hordei*-infected barley leaves at 4 and 7 dpi (Chen et al., 2019), enabling accurate identification of early-expressed effector genes. These assemblies represent the first high-contiguity phased reference for *P. hordei* and enable comparative analyses that were previously not possible due to genome fragmentation and allelic collapse (Chen et al., 2019). To identify candidate *Avr* genes in cereal rust pathogens, comparing the genome of a founding (parental) isolate with that of single-step mutationally-derived pathotypes exhibiting specific added virulence has been the most commonly used approach (Chen et al., 2017; Chen et al., 2019; Salcedo et al., 2017; Upadhyaya et al., 2021; Wu et al., 2020). The high-quality assemblies generated in the present study provide valuable resources for genetic studies of avirulence in *P. hordei*.

Our analyses revealed that *P. hordei* Chr9 exhibits features characteristic of core chromosomes that harbor accessory-like regions found in pathogens such as the southern corn leaf blight fungus *Cochliobolus heterostrophus* (Condon et al., 2013), or the accessory genomic regions in the vascular wilt causal agent *Verticillium dahliae* (Faino et al., 2016; Klosterman et al., 2011). Our data provides strong evidence to support the highly variable nature of Chr9 in both Ph560 and Ph518. Based on Hi-C mapping, in the Ph560 assembly there is no obvious phase switch in the contact map. This is evidenced by a clean diagonal and minimum off-diagonal noise when combining the two haplotypes, demonstrating predominantly *cis* contacts for both haplotypes and supporting correct phasing and a lack of assembly artefacts (Figure 1 and Supplementary Figure 1). Low similarity between Chromosomes 9A and 9B, large-scale haplotype-specific rearrangements, and reduced synteny between them further supported allelic structural divergence rather than phase switching. For the Ph518 assembly, despite a lack of Hi-C reads, the use of high-coverage HiFi reads resulted in highly continuous contigs. More than 30 contigs reached near-chromosome level assembly, with most haplotypes containing telomeric repeats including that of Chr9, similar to a previous report in *P. triticina* without using of Hi-C (Li et al., 2023). The same signatures of divergence between Chr9A and Chr9B seen in Ph560 were also observed in Ph518, which were isolated from the field at least 25 years apart. This independent recurrence is thus further strong evidence of biological reality, not assembly noise in Ph518.

Canonical accessory chromosomes such as those found in plant fungal pathogens like *Z. tritici* (Plissonneau et al., 2018), *Fusarium oxysporum* (Ma et al., 2010), and *Leptosphaeria maculans* (Grandaubert et al., 2014) are typically dispensable and lineage-specific. Chr9 in *P. hordei* is conserved in structure across both Ph560 and Ph518 but is highly enriched for signatures of structural plasticity and non-colinear regions (Figure 2A). This includes extensive translocations, duplications, and a large repeat-rich region lacking genes but flanked by putative effectors (Figure 2A, Figure 3, and Supplementary Figure 7). These features are further supported by Hi-C data that show compartmentalization breakpoints and spatial insulation at repeat-rich windows, particularly near the centromeric region (Figure 3 and Supplementary Figure 7). Consistent with the large-scale SVs in Ph560 (Figure 2A), compartment quantile analysis showed that Chr9A is not only compositionally distinct but is also organized into spatially alternating chromatin domains. This therefore indicates a unique structural and functional compartmentalization across haplotypes in *P. hordei*, represented by one particular chromosome pair instead of whole genome wide.

In particular, the elevated TE content and gene-sparse regions in Chr9A of Ph560 are consistent with the previous finding in *Verticillium dahlia* that TEs may drive diversification of accessory genomic regions by triggering large-scale rearrangements, duplication, and loss (Faino et al., 2016). Chromosome-specific structural variants have been linked to effector diversification and virulence adaptation in plant pathogens (Hartmann, 2022). Chr9 is enriched in simple repeat regions that are flanked by dense clusters of effector-like genes, suggesting it may act as a key genomic region for adaptive evolution in response to selective pressures. Notably, Chr9A uniquely exhibits a polarity inversion between gene density and TE content compared to the rest of the genome, with gene-rich, TE-poor compartments corresponding to low E1 domains, and TE-rich, gene-poor compartments falling into high E1 domains (Figure 3 and Supplementary Figure 7). This compartmental inversion was absent in Chr9B, where TE and gene distributions did not follow the same spatial partitioning, indicating strong haplotype-specific chromatin organization. The large-scale SVs in Chr9A are mainly at sub-telomeric ends, in line with regions known in other plant pathogenic fungi to be hotspots for genome rearrangement (Frantzeskakis et al., 2019; Grandaubert et al., 2014; Plissonneau et al., 2018). Unlike previously reported genomes of cereal rust pathogens including *Pt, Pgt,* and *Pst*, in which no chromosomes with accessory-like plasticity have been characterized, *P. hordei* Chr9 represents a novel example of a conserved chromosome enriched for structural variation and accessory features. These findings imply that Chr9 may function as a “compartmentalized” core chromosome with regions subject to rapid diversification, potentially playing a role in adaptive evolution of *P. hordei*.

The value of long-term continental-scale pathogenicity surveys and of historical collections of viable pathogenically defined rust isolates derived from them in understanding host adaptation and in resolving lineage-specific evolutionary patterns was highlighted by studies of Australian populations of the wheat stripe rust pathogen *Pst* (Ding et al., 2021). Population genomic analyses of *P. hordei* isolates representing pathotypes detected in Australia over a 54-year period (Luig, 1985; Cotterill et al., 1995; Park, 2003; Park RF unpublished) revealed five major genetic groups. Multiple complementary analyses including PCA analysis, ADMIXTURE, SplitsTree networks and f-branch statistics consistently revealed the presence of both clonally derived lineages and sexual and possibly asexual recombinant individuals. ADMIXTURE analysis recovered the same five population clusters across both haplotypes, showing high consistency in cluster assignment for clonal groups and revealing mixed ancestry in intermediate individuals. In particular, Group 2 and Group 5 formed well-defined clonal groups with minimal admixture, likely representing historical or recent asexual expansions within groups, whereas Group 1 and Group 3 isolates exhibited mixed ancestry, consistent with recombination and gene flow.

The co-ancestry matrices from both haplotype references were generally concordant but revealed subtle differences. Some individuals (e.g., Ph484, Ph506, Ph531) exhibited partial signals of both structure and admixture, suggesting either ancestral recombination or recent divergence followed by clonal spread. Across all datasets, Ph518, the most avirulent isolate, is clearly a basal genotype with minimal co-ancestry to any group, no evidence of admixture, and high divergence across PCA, ADMIXTURE, and SplitsTree results. These results collectively support the conclusion that *P. hordei* in this study has propagated via a mixed reproductive strategy in Australia for over 50 years. The presence of large clonal groups with minimal genetic differentiation suggests that asexual urediniospore propagation is common and effective over multiple seasons or regions. However, the detection of clear recombinant individuals, supported by ancestry painting, f-branch introgression signals, and reticulation in SplitsTree, indicates that sexual or parasexual reproduction has occurred either *via* the alternate host or through somatic hybridization. The variability in co-ancestry asymmetry and admixture proportions further suggests that these recombination events are not uniform but occur sporadically, potentially contributing to genetic diversity and virulence evolution.

We further provided clear evidence to support reproductive patterns across *Ph* population isolates. The two isolate groups from k-mer analysis of Chr9 are in line with the reference-based genome-wide variation data. The most variable k-mers from each chromosome produced concordant binary presence/absence profiles, with isolates partitioned into two groups. Collapsed k-mer clusters derived by sequence similarity highlighted the redundancy among these k-mers. Notably, the most divergent k-mer clusters are enriched for an AAACAT-type motif often found in Forkhead transcription factor binding sites that are known in yeast to regulate cell-cycle timing and mating-type switching (Sun et al., 2002). These motifs are likely located in or near cis-regulatory elements of the mating-type locus, and their divergence supports a model in which chromosomal haplotypes encompassing these motifs have co-evolved with reproductive identity. The k-mer and mating-type gene analyses not only confirmed the SNP-based population structure but also explained its mechanistic basis. Reproductive isolation appears to be maintained by fixed mating-type haplotypes, and divergence at these loci is predictive of broader genomic divergence. This mirrors observations in other rust fungi, where sexual lineages exhibit mating-type polymorphism and recombination, while asexual lineages carry non-recombining, fixed mating-type haplotypes (Holden et al., 2023; Luo et al., 2024). In *P. hordei*, the k-mer signature of the mating-type chromosomes is both a marker of and a likely contributor to the evolutionary separation between sexual and asexual lineages.

We also demonstrate, for the first time, that CNV is a primary driver of population-level divergence in *P. hordei*. CNV profiling across 40 *P. hordei* isolates revealed clear temporal trajectories, with recent isolates particularly those from the widespread 5453 P clonal lineage accumulating extensive gene duplications (Figures 6A-C). These duplications were strongly enriched on Chr9A for the Group 5 isolates (Supplementary Figure 13A), supporting its role as a structural hotspot for adaptive evolution. Moreover, we showed that CNV patterns correlate tightly with clonal groupings (Figures 6A-C and Supplementary Figures 13A-B), suggesting that structural variants are not only the consequences of, but may also act as drivers of, population shifts under selection.

Our study provides the first functional evidence linking CNV to fungicide insensitivity in a rust fungus. We identified predominant duplications of *Cyp51* in isolates from three fungicide insensitive isolates that belong to the 5453 P lineage, which exhibited high levels of fungicide insensitivity in phenotypic assessments (Park et al., 2022). Detailed fungicide sensitive and insensitive categorization are ongoing to assess the prevalence and evolutionary dynamics of CNV across broader *P. hordei* populations and other rust pathogens. Alleles of *Cyp51* are exactly Chr9 SV-hotspot localized, consistent with our CNV findings and verified via qPCR. By mapping Ph560 reads to the references of itself, Ph560 naturally served as a CNV threshold standard, thus accurately reflecting global CNV differences across all isolates. *Cyp51* encodes sterol 14α-demethylase, an enzyme that is inhibited by DMI/triazole fungicides in many fungal pathogens, and in which mutations can lead to DMI insensitivity (Cook et al., 2021; Ma et al., 2006; Rosam et al., 2021). Functional assays in a *S. cerevisiae erg11* mutant confirmed that the *P. hordei Cyp51* allele restores azole insensitivity, directly linking CNV to fungicide tolerance. This is distinct from the prior genome-wide studies in *Pst* (Cook et al., 2021), where *Cyp51* sequence variation was observed but lacked phenotypic validation. Our data thus established an essential link between structural genome variation and fungicide resistance evolution in rust fungi. This observation is consistent with CNV-associated fungicide resistance in the barley pathogen *R. commune*, where segmental duplications spanning *CYP51A* contribute to reduced azole sensitivity (Stalder et al., 2023). CNVs were proposed as a major source of adaptive genetic variation for recent adaptation of *R. commune* (Stalder et al., 2023). While similar adaptive mechanisms have been reported in human fungal pathogens such as *C. albicans* and *Cryptococcus neoformans*, *P. hordei* represents the first rust species where *Cyp51* CNVs have been detected and functionally validated. Using haplotype-phased Ph560 assemblies and comparative read-depth profiling between the two haplotypes, we resolved duplications at the *Cyp51* locus on Chr9 with high confidence, providing key insight into genomic architecture of fungicide resistance.

## Acknowledgements

Financial support from Judith and David Coffey and family, the Grains Research and Development Corporation (GRDC: US221; US315; US00067; UOS1707-003RTX), and the University of Sydney is gratefully acknowledged. This research was undertaken as part of a long running program on national cereal rust surveillance, conducted at the University of Sydney since 1921. We would like to acknowledge technical assistance provided by Ashley Grewar, Margerita Pietilainen, and Matthew Williams in purifying and maintaining rust isolates.

## Author contributions statement

Y.D. and R.F.P. designed this study. Y.D. and X.Y. conducted the analyses and drafted the manuscript. R.F.P., P.Z., M.C., T.R., X.Y., and Y.D. revised the manuscript. X.Y., P.Z., M.C., and Y.D. performed the DNA experiments. M.H. prepared materials for Hi-C sequencing. All authors approved the submitted manuscript and declare no conflict of interest.

## Notes

### Competing Interest Statement

The authors have declared no competing interest.

