## Supplementary Figure for "Genome structural plasticity and copy number dynamics shape adaptive evolution in a barley fungal pathogen"

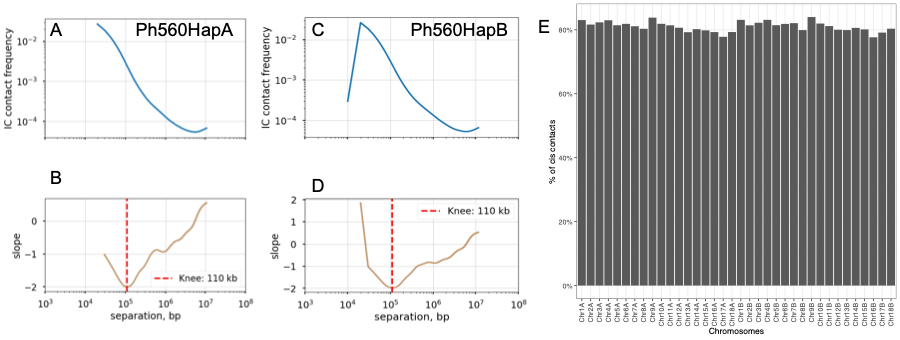

**Supplementary Figure 1:** HiC contacts assessment for the Ph560 reference genome assembly. (A-D) (E) Percentage of cis-contacts in all 36 chromosomes indicating minimalized phase switch.

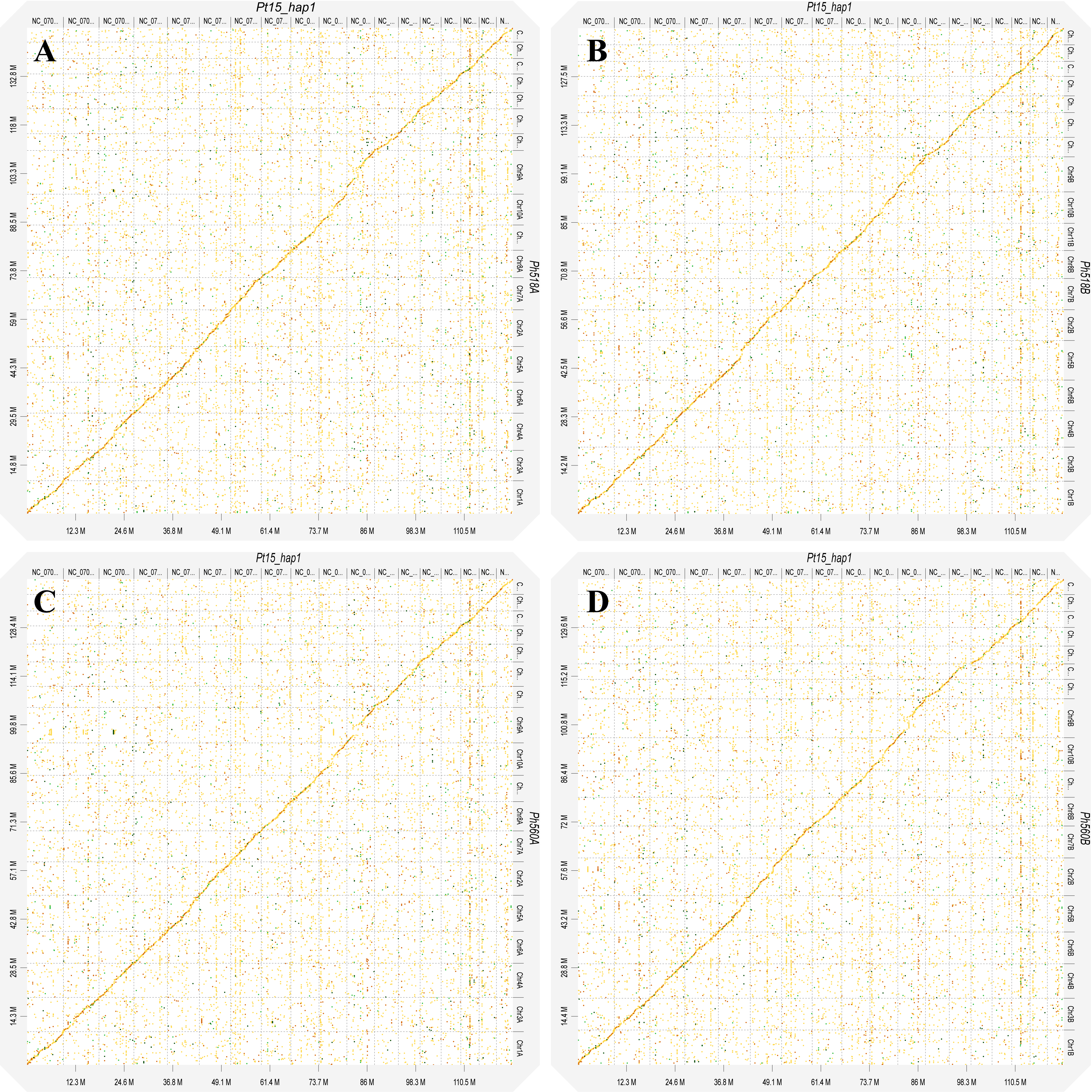

**Supplementary Figure 2:** Genome alignment between a chromosome-level *Puccinia triticina* reference (Pt15 haplotype 1) and the four haplotypes (queries) of *Puccinia hordei*, demonstrating clear synteny across the genome for all four haplotypes. (A): Ph518A, (B): Ph518B, (C): Ph560A, and (D): Ph560B.

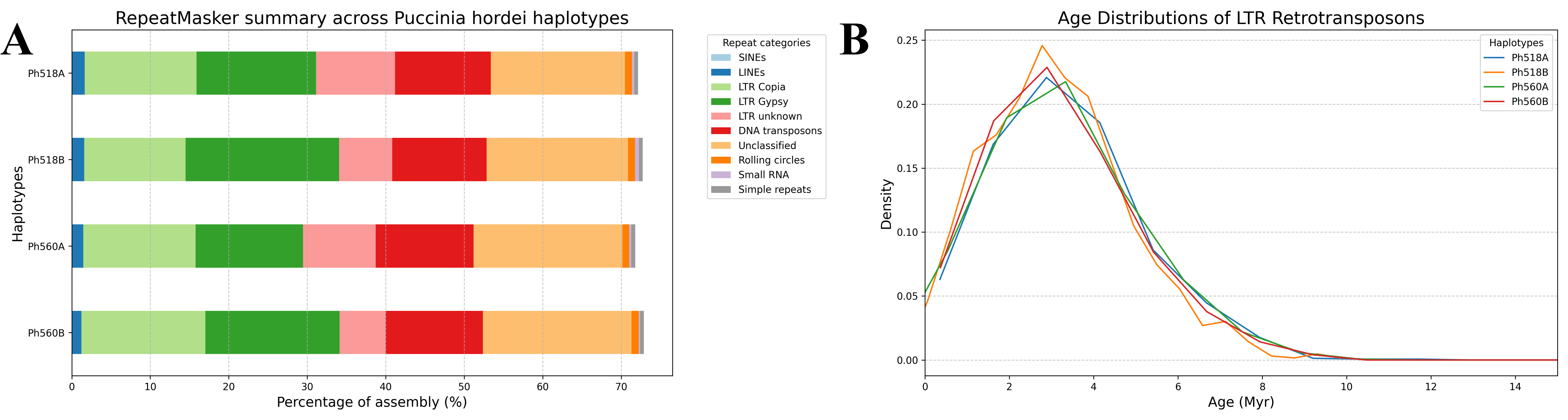

**Supplementary Figure 3:** (A) Summary of repeat content in each *Puccinia hordei* haplotype, annotated by RepeatMasker. Colors represent different repeat categories. (B) Distributions of estimated insertion times for LTR-retrotransposons over evolutionary history, calculated using a substitution rate of 2% per million years. A burst of LTR-retrotransposon expansion is observed at approximately 3 million years ago across all four *Puccinia hordei* haplotypes.

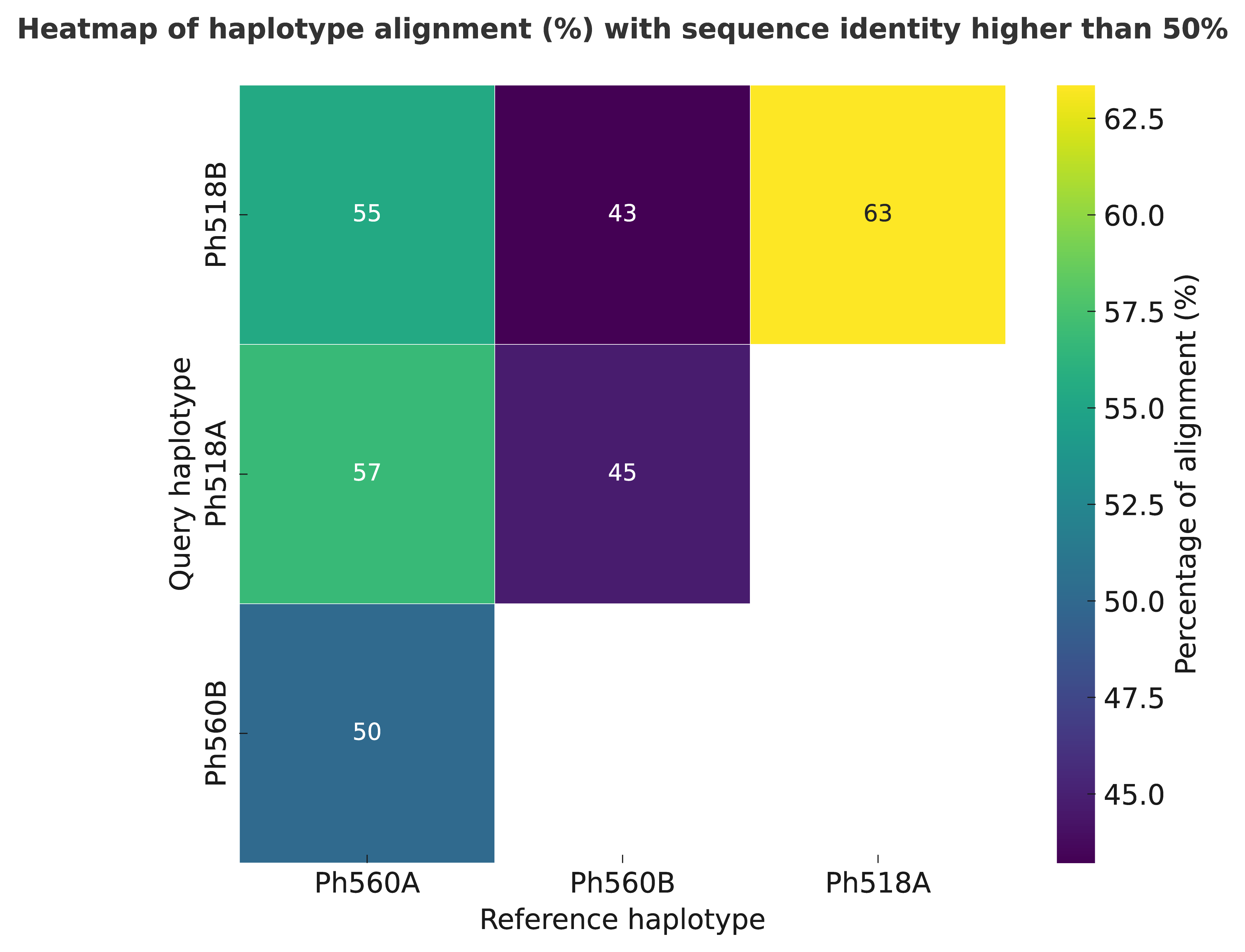

**Supplementary Figure 4:** Heatmap showing the percentage of alignment with sequence identities greater than 50% between *Puccinia hordei* haplotypes, revealing that Ph518A and Ph518B are the most similar, while Ph518B and Ph560B are the most divergent haplotypes in terms of sequence similarity. Haplotype alignments were performed pairwise using D-Genies.

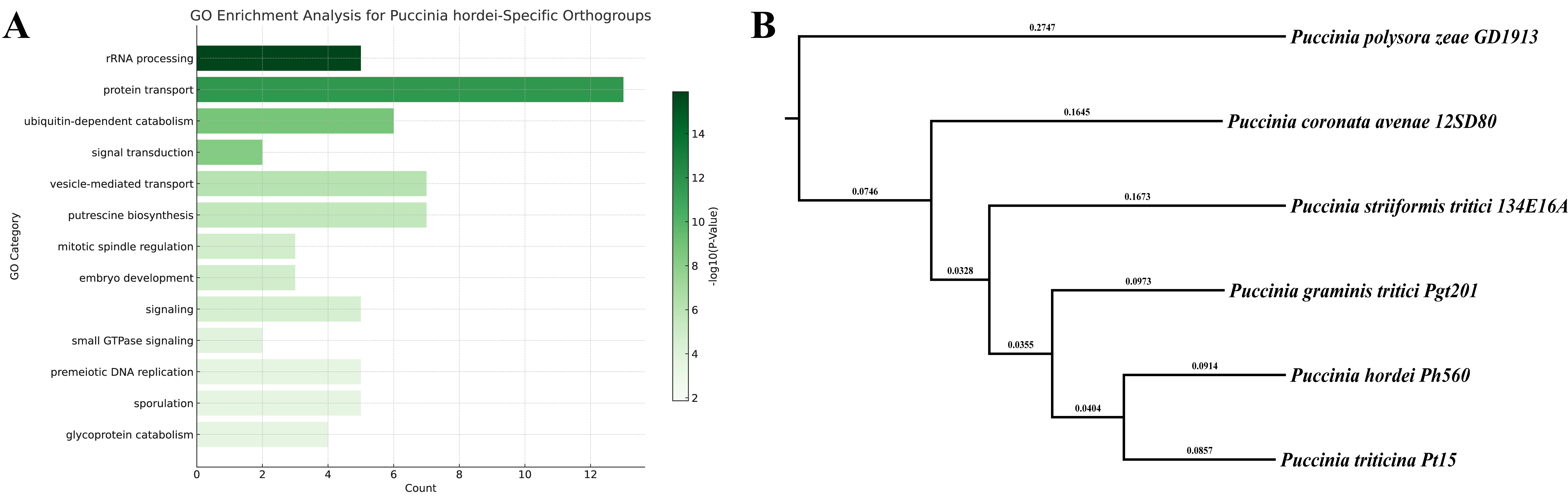

**Supplementary Figure 5:** (A) GO enrichment analysis of *Puccinia hordei*-specific orthogroup proteins compared to orthogroups from five other *Puccinia* species. (B) Maximum likelihood phylogenetic tree of six *Puccinia* species based on single-copy orthogroup proteins, highlighting that *Puccinia hordei* and *Puccinia triticina* are the most closely related species, despite differences in host specificity.

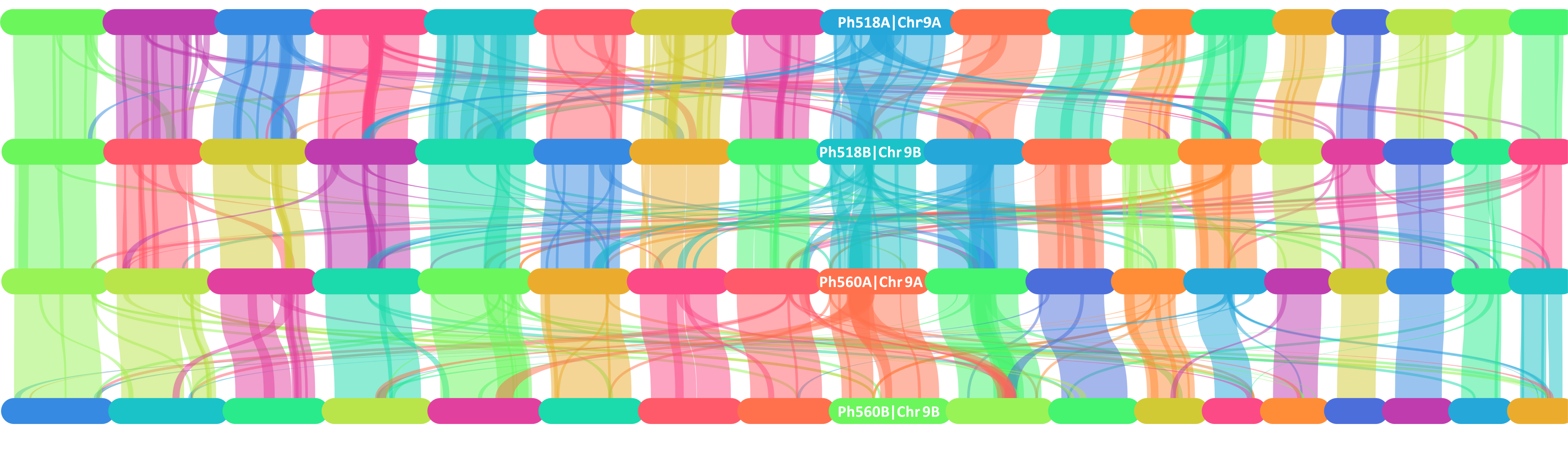

**Supplementary Figure 6:** Collinearity analysis between the four *Puccinia hordei* haplotypes. Extensive synteny was observed across the haplotypes for most of each genome. Disrupted or shuffled regions reflect small structural variations.

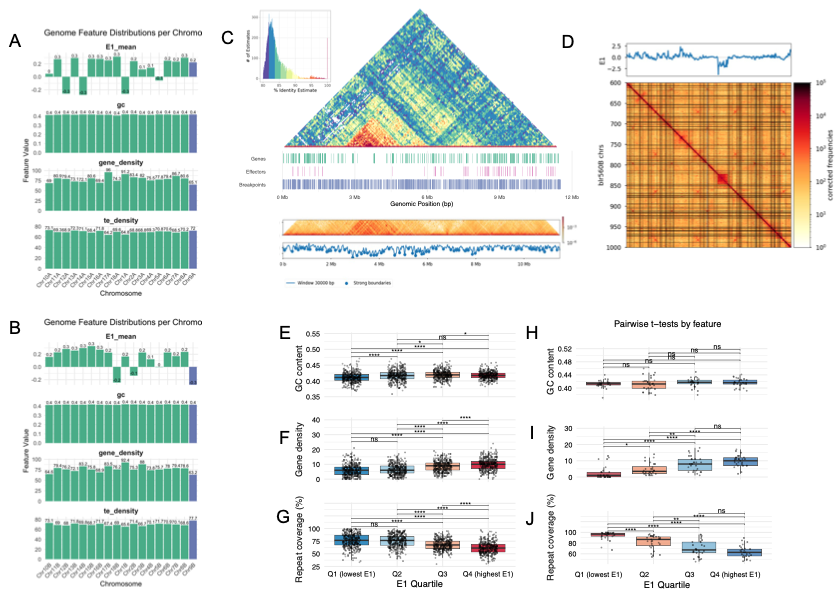

**Supplementary Figure 7:** Overview of Ph560 genome feature distributions per chromosome and Chr9B characteristics. Mean first eigenvector (E1) values, GC contents, gene and TE density ratio were calculated for all chromosomes for both hapA (A) and hapB (B). Chromosomes 9 were highlighted in blue. (C) Whole-chromosome self-alignment of Chr9B. Gene and predicted effector annotations (green and purple ticks) and inter-isolate alignment between Ph560 Chr9A and Ph518 Chr9A are shown along the axes. SV coordinates between Ph560 Chr9B and Ph518 Chr9B were represented by breakpoints. The upper display indicates percentage identity distribution of alignments. Lower panel heatmap showing Hi-C contacts map across Chr9B. Topology associated domains in 30kb window were predicted and indicated by blue dots as strong boundaries. (D) Hi-C contact matrix and first eigenvector (E1) of Chr9B in Ph560. Gids show compartment-like domains spanning bins 600-1000 (chromosomes 7-10). Heatmap indicates corrected contact frequencies. Crosses reflect centromere locations. (E-J) Genomic features in compartments of the Ph560 hapA genome. (E-G) Genome-wide distributions of GC content, gene density (number of genes), and TE coverage across Hi-C E1 quartiles. Q1 (lowest E1) is associated with low GC/gene content and high TE load, while Q4 (highest E1) shows the opposite. (H-J) Chr9B compartmental distributions of the same genomic features.

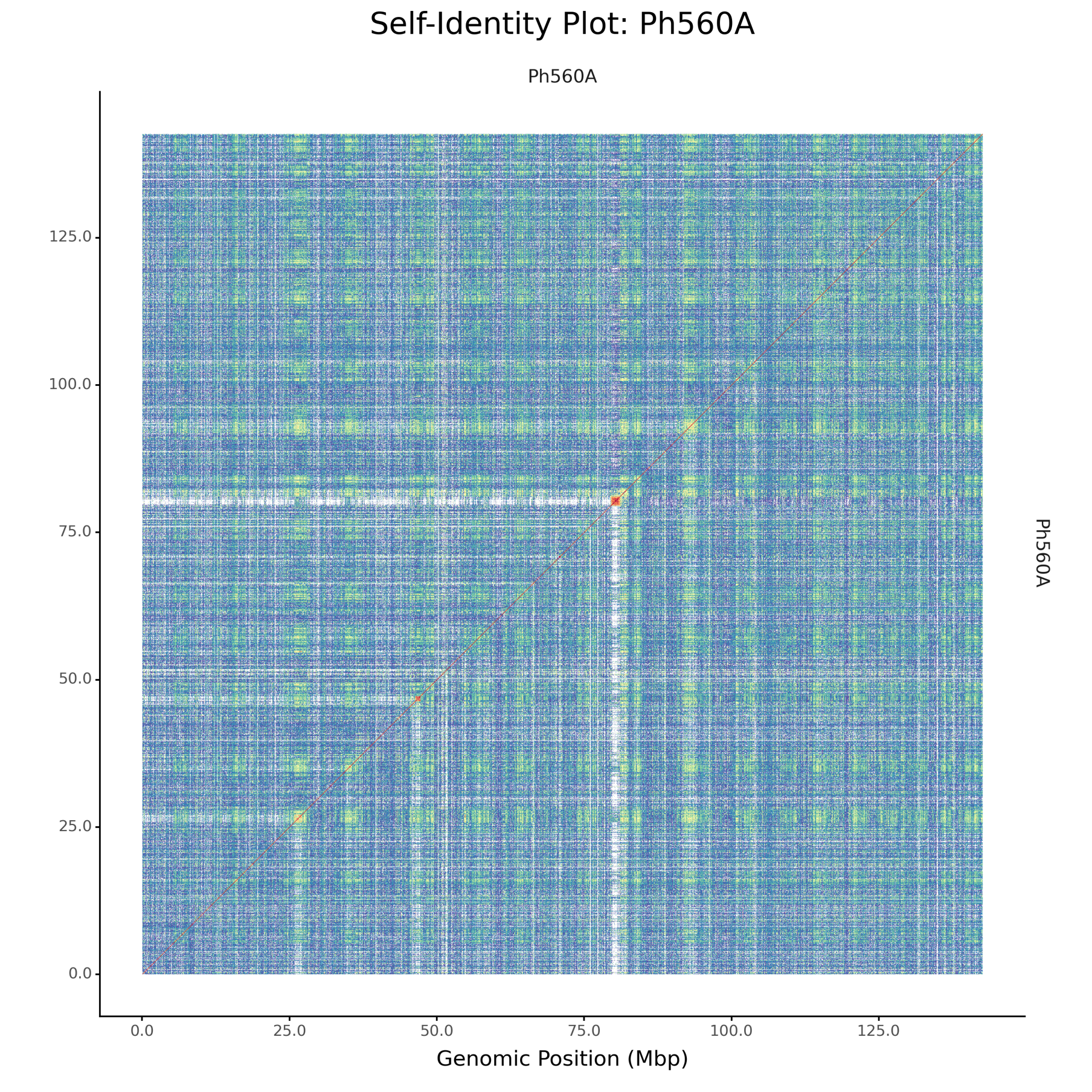

**Supplementary Figure 8:** Whole-genome alignment of Ph560 haplotype A.

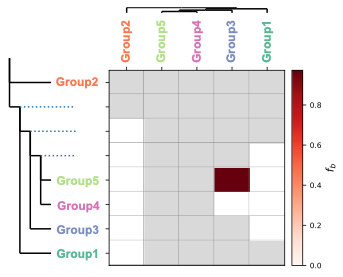

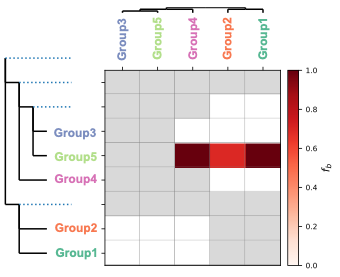

A

B

C

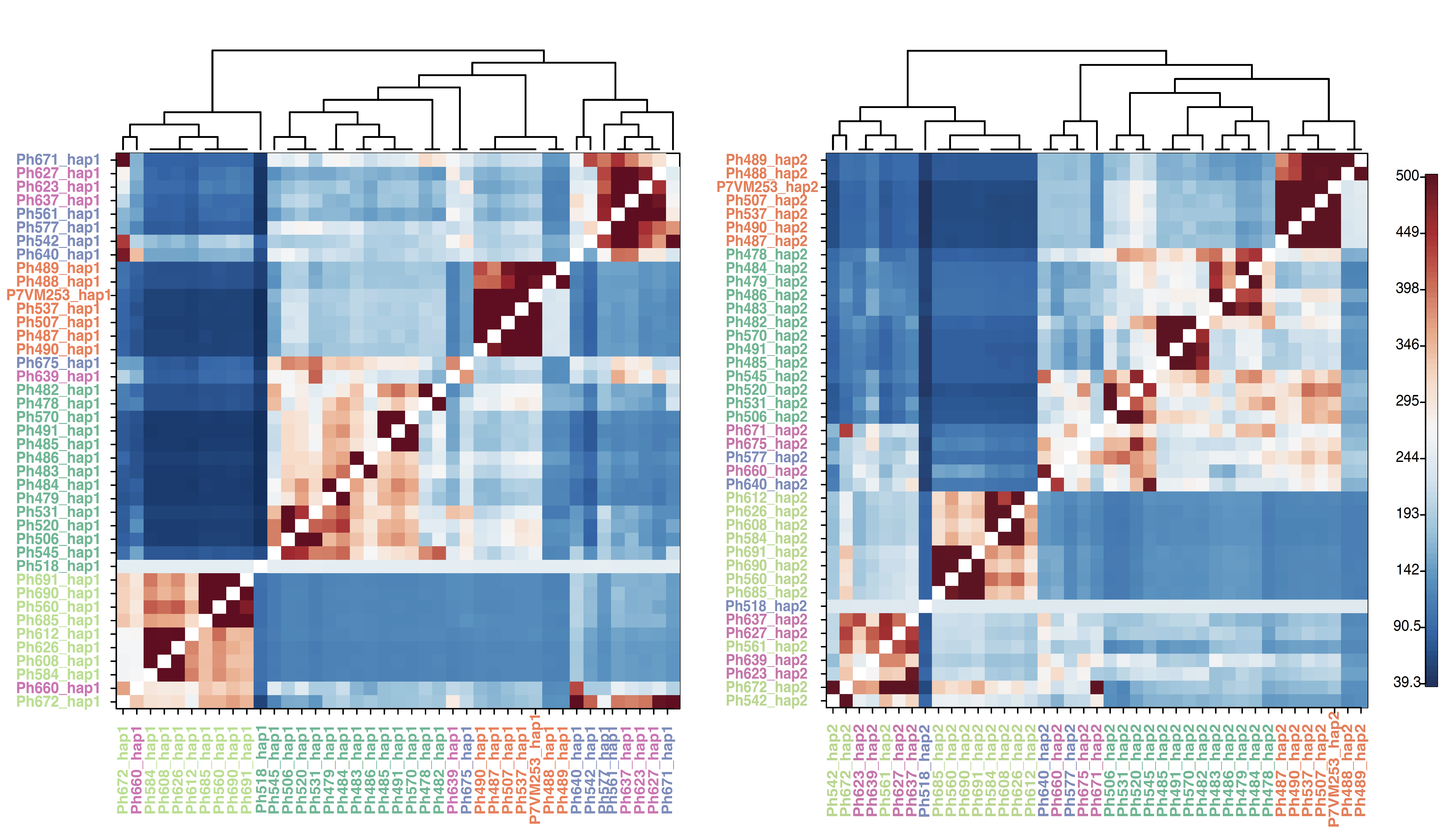

D

**Supplementary Figure 9:** F-branch test and co-ancestry analyses. F4-ratio statistics for all possible trios of five Groups defined by PCA analysis. Heatmaps show the f-branch statistic (color scale) computed by Dsuite, summarizing excess allele sharing between each internal branch of the species tree (rows) and each tip Group(columns), relative to the tree topology. Top tree was based on phylogenetic relationships of all isolates in hapA and hapB represented by Groups. Each internal branch (row) is tested by aggregating D‑statistics across quartets that contrast its descendant clade with the rest, asking whether the target population shares more alleles with that internal branch than expected. (A) Results based on the hapA reference genome. (C) Results based on the hapB reference genome. Dark red cells indicate stronger evidence of gene flow or shared ancestry between the corresponding branch and Group. (B and D) Pairwise coancestry matrices among individuals inferred from haplotype sharing. Coancestry heatmaps based on hapA (B) and hapB (D) reference genomes. Each cell represents the number of DNA chunks donated (rows) and received (columns) between individual pairs, with darker red colors indicating higher sharing. Individuals are clustered hierarchically based on shared ancestry, and color‑coded by population group. Clustering (top) is based on maximum likelihood ordering with the default fineSTRUCTURE tree building algorithm.

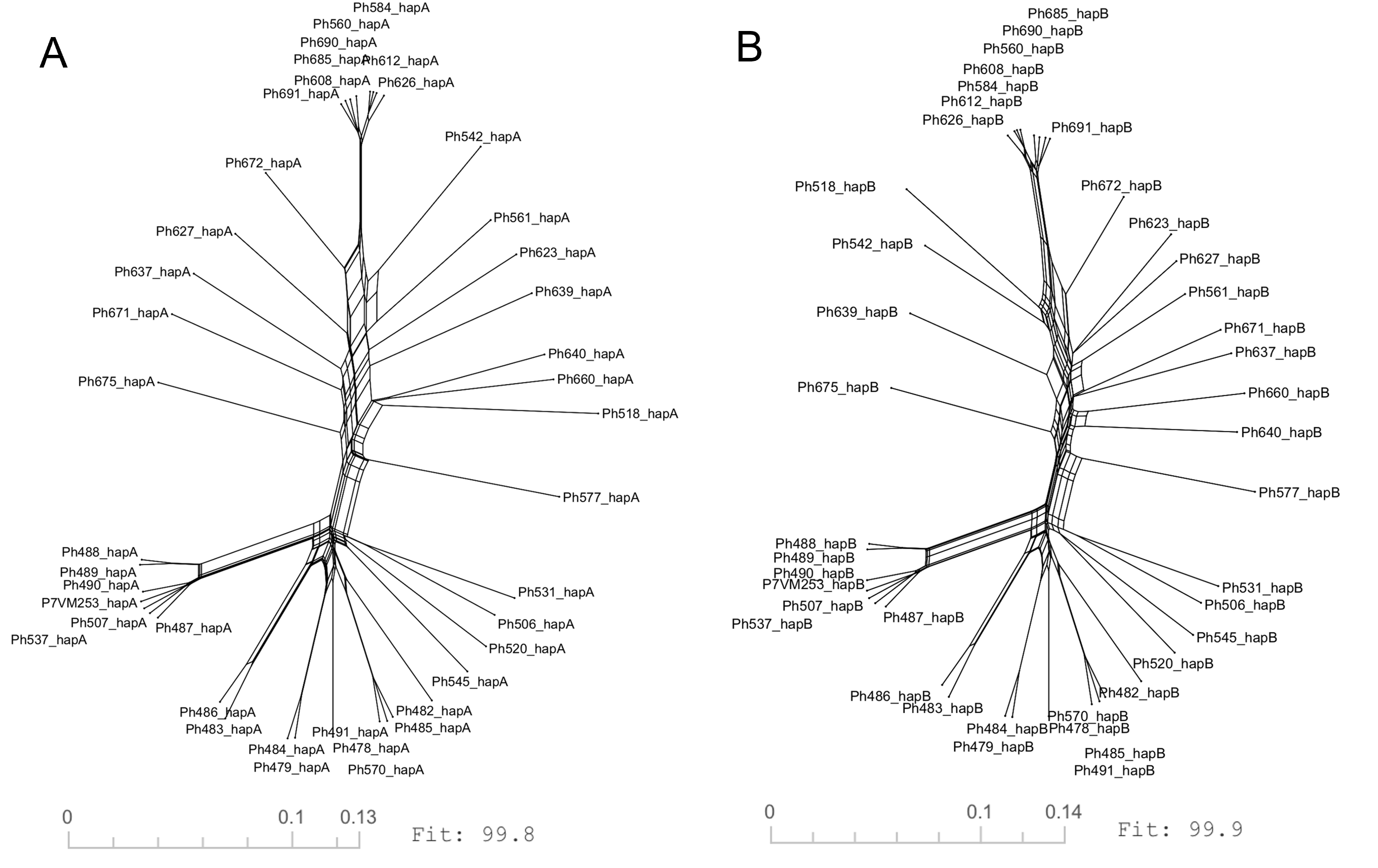

**Supplementary Figure 10:** Split network trees representing genetic relationships among *Ph* individuals based on hapA (A) and hapB (B) reference genomes. The networks reveal clusters of closely related individuals as well as signals of admixture or recombination, shown as parallel edges or network-like structures. Fit values indicate the proportion of variance explained by the network.

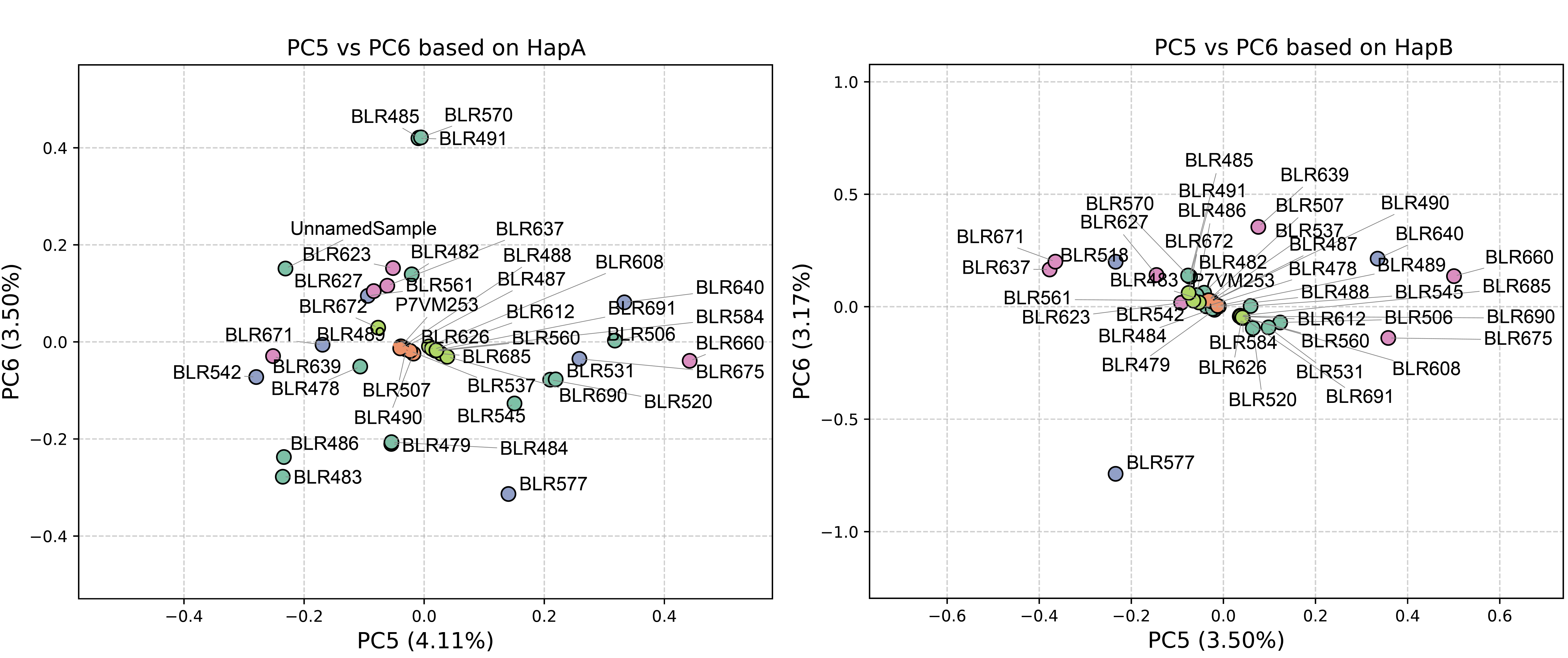

**Supplementary Figure 11:** PCA analysis major axes of variation between PC5 and PC6.

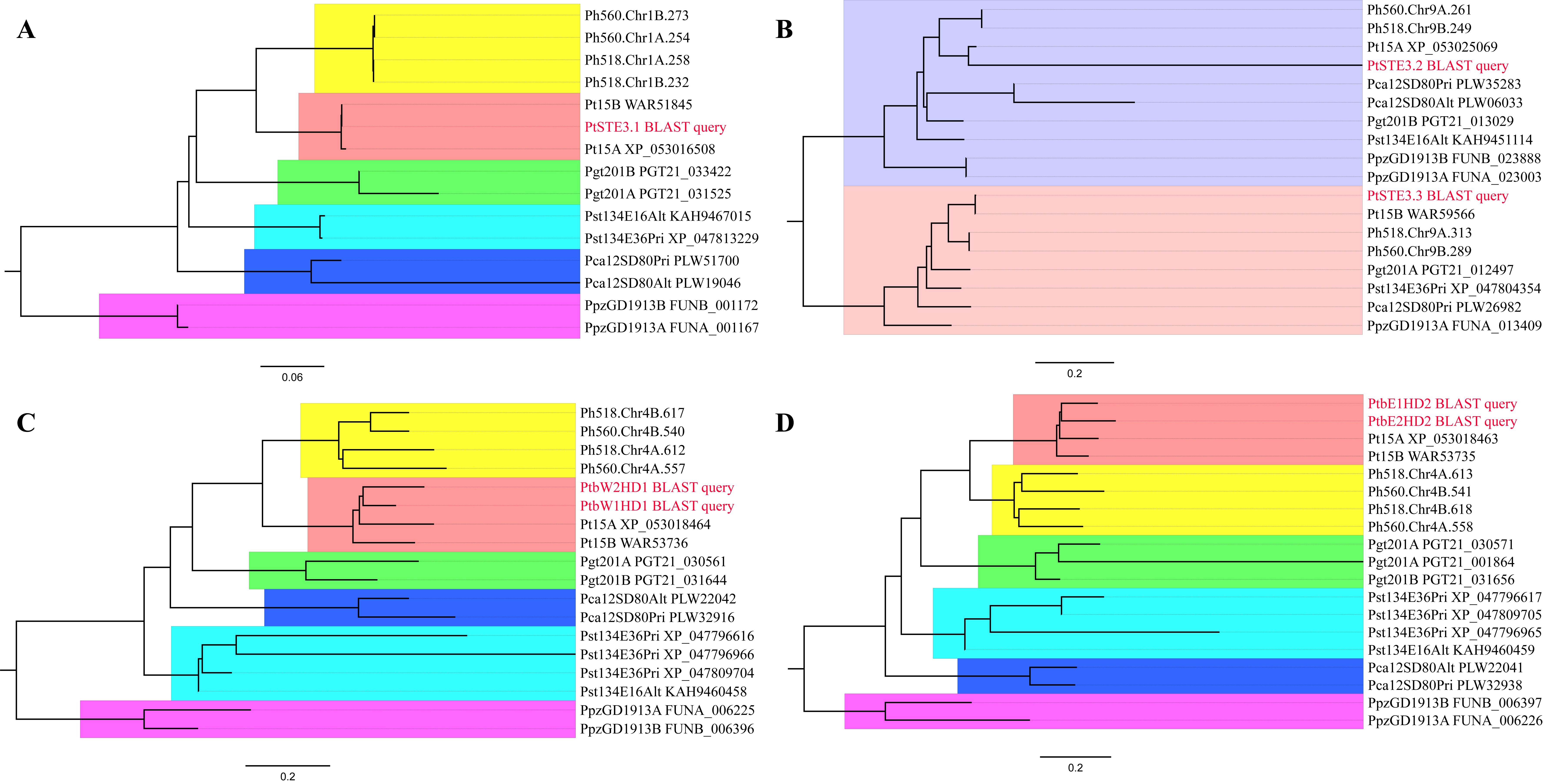

**Supplementary Figure 12:** Clusters of mating type genes classified by orthogroup analysis using OrthoFinder3: *STE3.1* (A), *STE3.2* and *STE3.3* (B), *bWHD1* (C) and *bEHD2* (D). The queries used for sequence blasting are colored in red. Clades are marked by different colors.

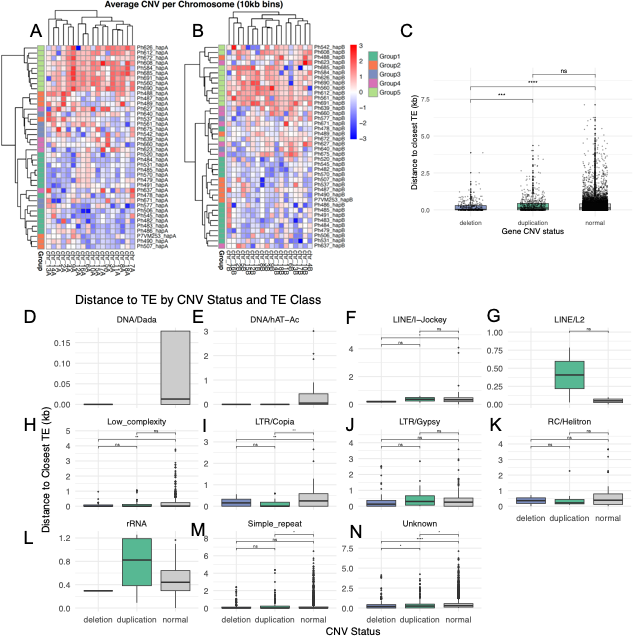

**Supplementary Figure 13:** CNV distibutions in the Ph560 reference genome. (A-B) Average number of CNVs per chromosome in hapA and hapB genomes in a 10kb window. Heatmap showing log2 transformed CNV numbers. (C) Distance of genes showing deletion, duplication or normal status across all *Ph* samples to the nearest TE. (D-N) Distance of each gene types to specific TE element classes. Student’s t-test: **p*<0.5 ***p*<0.01; *** *p*< 0.005; **** *p*<0.001.

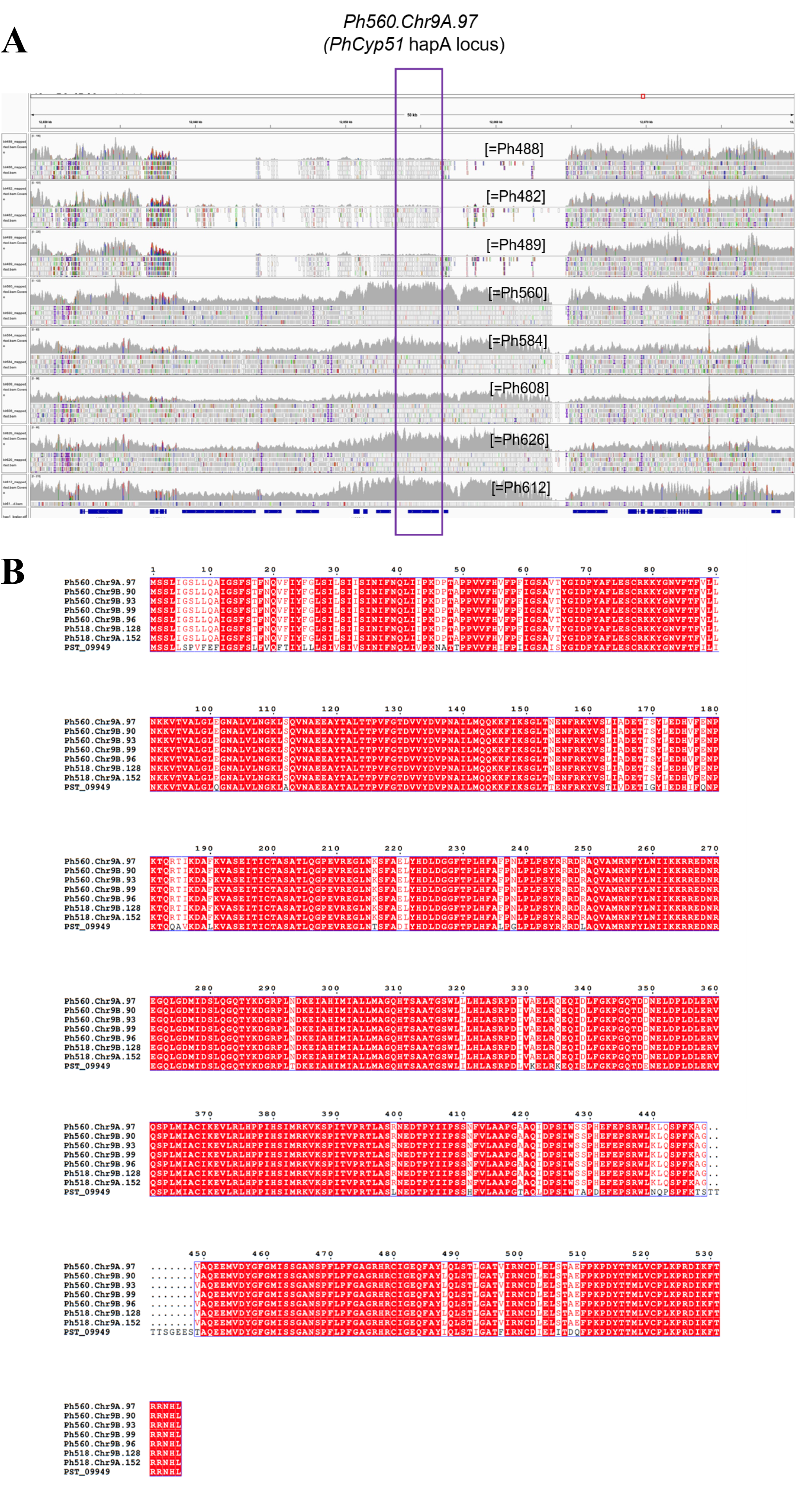
**Supplementary Figure 14:** (A) Representative IGV overview of reads alignment in the hapA genomic window containing the Cyp51 locus. (B) Protein sequence alignment of all homologs of PhCYP51 in Ph560 and Ph518 showing no sequence variation.
